# Behaviorally gated hierarchical predictive coding in macaque auditory–prefrontal circuits

**DOI:** 10.1101/2025.08.10.669259

**Authors:** Haoxuan Xu, Peirun Song, Hangting Ye, Ana Belen Lao-Rodriguez, Qichen Zhang, Yuying Zhai, Xuhui Bao, Ishrat Mehmood, Hisashi Tanigawa, Zhiyi Tu, Lingling Zhang, Xuan Zhao, David Pérez-González, Manuel S. Malmierca, Xiongjie Yu

**Author notes:** Corresponding authors (X.Y.), (M.S.M). These authors contributed equally (Haoxuan Xu, Peirun Song, Hangting Ye, Ana Belen Lao-Rodriguez). Lead contact: Xiongjie Yu. Author contributions: Conceptualization, X.Y.; Methodology, X.Y., H.X. and P.S.; Investigation and Data Curation, H.X., P.S., H.Y., Q.Z. and Y.Z.; Writing-Original Draft, X.Y., H.X., P.S., A.B.L.R., and M.S.M; Writing-Review & Editing, H.X., P.S., X.Y., A.B.L.R., D.P.G. and M.S.M; Funding Acquisition, X.Y., Y.Z. and M.S.M; Resources, H.T. and X.Y.; Software, H.X., P.S., H.Y., X.B., Y.Z., I.M., Z.T., L.Z. and X.Z.; Validation and Visualization, H.X. and P.S.; Supervision, X.Y., M.S.M. Declaration of interests: The authors declare no competing interests.

## Abstract

The ability to detect deviations from expected sensory input is fundamental for adaptive behavior. We recorded electrocorticographic activity from the auditory (AC) and prefrontal (PFC) cortices of behaving macaques during an auditory oddball task to probe the cortical dynamics of predictive processing. Repetition of standard stimuli evoked suppression and facilitation in AC and strong low-frequency (2 Hz) enhancement in PFC, accompanied by bidirectional delta-band coupling indicative of a shared predictive state. Deviant stimuli triggered early AC responses followed by PFC activation and increased feedforward and feedback connectivity across delta, theta, and gamma bands. Behavioral engagement amplified both prediction and prediction error signals, strengthening cortical network coordination. Together, these findings reveal a hierarchical predictive network in which the AC encodes sensory regularities and violations, while the PFC integrates predictive context in a behaviorally dependent manner.

## Introduction

Living organisms are continuously exposed to a storm of sensory stimuli, yet only a small fraction is behaviorally meaningful, while most are routine and can be disregarded. Detecting changes or novel events is therefore essential for adaptive behavior, and sensory salience depends strongly on context and prior experience^1–5^. To handle this continuous stream of input efficiently, the brain relies on a proactive computational framework known as predictive coding, which uses prior expectations to guide the processing of incoming sensory information. The predictive coding theory posits that the brain continuously generates active predictions (top-down processing) based on sensory signals from the external environment^6–8^. These predictions are then compared to actual sensory input (bottom-up processing), producing prediction errors that serve as signals to update the system.

The auditory system is particularly efficient at filtering complex acoustic input into manageable representations^9–11^. At the neuronal level, some auditory neurons reduce their firing to repeated sounds while maintaining sensitivity to novel ones, a phenomenon known as stimulus-specific adaptation (SSA)^12–25^. At the macroscopic level, mismatch negativity (MMN), an event-related potential observed in human electroencephalogram (EEG), reflects a similar mechanism of auditory deviance detection^26–30^. SSA and MMN have been extensively investigated using the classical oddball paradigm in animal models, with evidence of their presence across multiple brain regions (including the auditory midbrain and cortex) and across species and arousal states^12–14,16,29,31–34^. Despite these advances, most animal studies have employed acoustically simple, behaviorally irrelevant sounds, leaving key gaps in understanding how prior experience shapes SSA and MMN or how these responses reflect neuronal plasticity^11,19,29,35–37^. Moreover, many findings come from passive conditions in which subjects were not engaged in cognitive or behavioral tasks, highlighting the need to explore SSA and MMN within meaningful behavioral contexts^19,29,38,39^. Non-human primates, whose auditory and cognitive systems closely resemble those of humans, offer an ideal model for such research^40–45^. Macaques, in particular, display human-like auditory abilities including speech discrimination, sound localization and sensitivity to surprising sounds, making them especially suited for investigating the neural mechanisms of change or novelty detection^41,46–48^.

SSA and MMN represent neural correlates of prediction errors that intensify along the auditory hierarchy, from the inferior colliculus (IC) to the auditory cortex (AC), and from primary (lemniscal) to higher-order (non-lemniscal) regions^23,34,49,50^. These findings underscore the hierarchical nature of novelty processing but provide limited insight into the interregional, long-range cortico-cortical dynamics underlying prediction and prediction error. It has been proposed that predictions and errors operate in distinct frequency bands—low (alpha/beta oscillations) frequencies for predictions and high (gamma oscillations) frequencies for errors—originating from higher and lower cortical areas, respectively^51–57^. However, direct experimental evidence remains elusive. Testing these hypotheses requires simultaneous recordings across multiple levels of the hierarchy. The AC and prefrontal cortex (PFC) are particularly suitable regions to examine, as both play essential roles in novelty detection yet occupy distinct positions within the cortical hierarchy^31,50,56–60^. Importantly, the PFC is thought to contribute directly to the generation of predictive components^58–62^, highlighting the need to determine how top-down signals from the PFC are propagated and integrated with bottom-up inputs from sensory areas. Resolving this interaction remains a central challenge for understanding the neural circuitry that supports the predictive coding framework. Reciprocal anatomical connections between these regions have long been established^63^. Because the MMN typically emerges around 200 ms after a deviant tone, it has been interpreted as a prediction-error response derived from comparing the deviant input with predictions formed from the repeating standards^8^. Human EEG and electrocorticography (ECoG) studies^64,65^ show that a network linking the superior temporal gyrus in the temporal lobe with frontal regions is involved in MMN generation. Evidence from non-human primates further supports the view that the prediction-error components underlying MMN are distributed across the temporal and frontal cortices^66–69^.

Yet the respective roles of the PFC and AC, and their interactions during predictive processing, remain elusive. These mechanisms are fundamental for adaptive behavior, enabling organisms to anticipate events, detect discrepancies, and update expectations. Monkey ECoG studies have shown that top-down and bottom-up signals propagate through distinct frequency bands^70^, and assessing the frequency dependence of phase synchrony between the frontotemporal cortices can reveal the functional coupling that supports prediction and prediction-error responses. ECoG recordings from electrodes implanted in the temporal, lateral prefrontal, and orbitofrontal cortices of macaque monkeys have demonstrated synchronous low-frequency oscillations between frontotemporal regions during both tone presentation and omission^71^. Examining AC–PFC communication, particularly with directional metrics such as Granger causality (GC)^53,61,62^, may therefore clarify the circuitry underlying prediction and prediction error. Understanding these processes in behaving non-human primates is essential for advancing our knowledge of cognitive prediction mechanisms.

In this study, we investigated how the auditory and prefrontal cortices jointly implement predictive processing during active listening. Behaving macaques performed a behaviorally relevant auditory oddball task while we recorded electrocorticography activity simultaneously from both regions. Granger causality analyses revealed the directionality and frequency structure of interareal communication underlying prediction and prediction-error signaling. The two regions showed distinct repetition-dependent dynamics, enhanced low-frequency coupling during auditory regularity, and temporally precise prediction-error responses to deviant sounds. Critically, task engagement strongly amplified these effects, indicating that behavioral relevance sharpens both predictive and error-related representations. Together, these findings uncover a dynamic interplay between auditory and prefrontal cortices that supports context-dependent predictive coding and flexible auditory perception.

## Results

### ‘Prediction’ mechanisms during novelty detection in AC and PFC under behavioral conditions

To probe the neural and behavioral mechanisms of novelty detection, we trained two macaques to perform an auditory oddball task composed of sequential blocks of stimuli (Fig. 1A). Each block contained 7–10 repetitions of complex sounds with a 245-Hz fundamental frequency and seven harmonics (Fig. 1A bottom, see Methods for details). Each block ended with either a standard stimulus, identical to preceding ones, or a frequency deviant (Fig. 1A). Monkeys were required to press a button within 600 ms after deviants (hit trials) and to withhold responses to standards (correct rejections). A separate passive session was conducted after the behavioral session, during which the same oddball stimuli were presented without task engagement (Fig. 1B). Behavioral accuracy increased with the magnitude of frequency contrast or deviation (Fig. 1C). Neural activity was simultaneously recorded from two 64-channel ECoG arrays implanted over the AC and PFC cortices (Fig. 1D; Supplementary Fig. 1 shows electrode coverage and anatomical landmarks, including the superior temporal sulcus and lateral sulcus).

**Figure 1.**
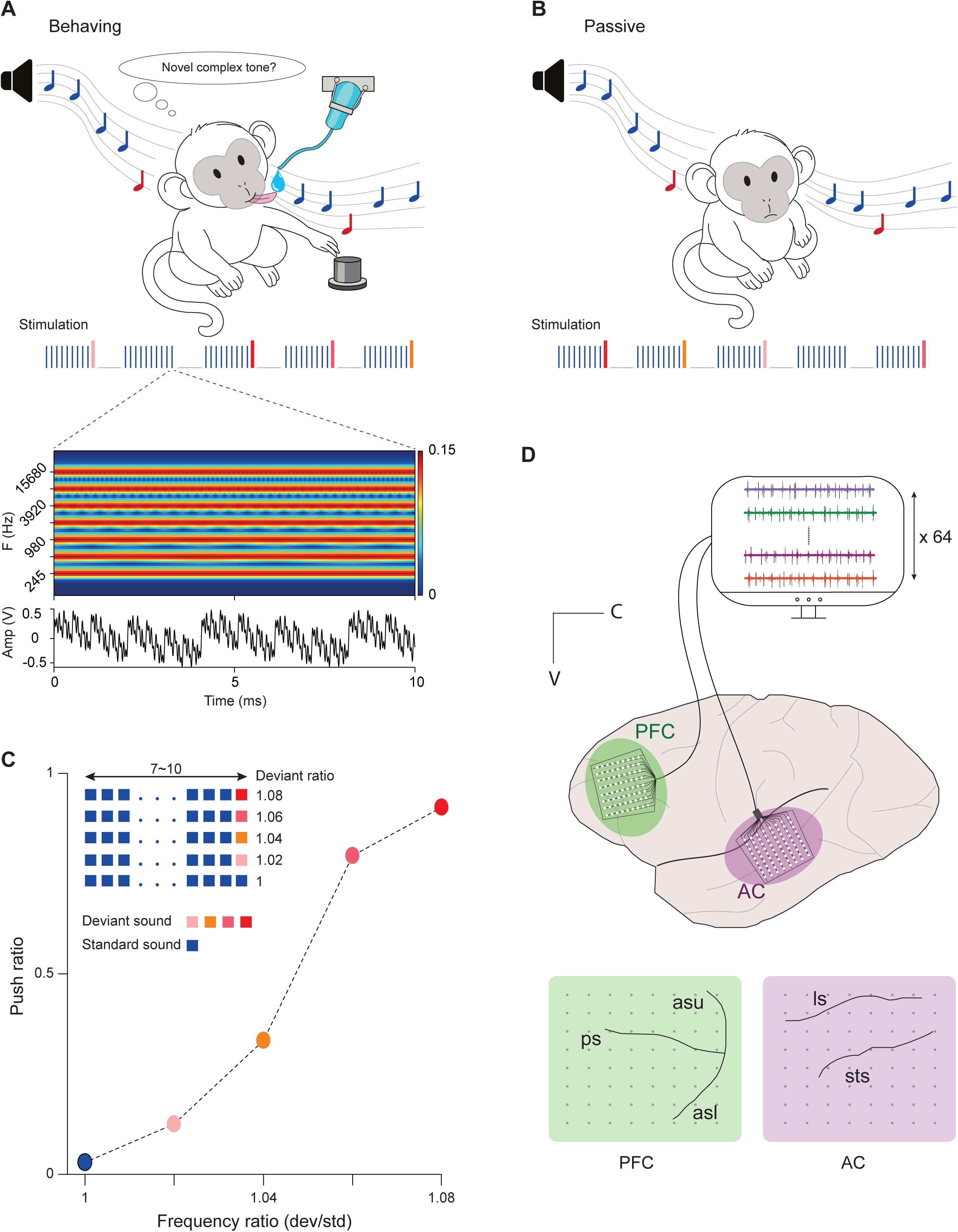
Experimental setup and task design for novelty detection in rhesus monkeys. **(A)** Auditory oddball task during the behaving session. Stimuli were presented in blocks of 7–10 repetitions of a standard complex tone (245 Hz fundamental and six additional octave-spaced harmonic components; bottom panel: 10-ms magnified view of stimulus spectrogram and waveform) followed by either the same tone or a frequency deviant. Deviants differed from the standard by frequency ratios of 1 (control), 1.02, 1.04, 1.06, or 1.08 (blocks with different deviants were presented randomly). Each complex tone was 100 ms in duration and was presented with a 500-ms inter-stimulus interval. Monkeys reported deviants by button pressing within 600 ms after deviant onset to obtain a reward. **(B)** Passive session in which the identical oddball stimulus sequences were presented without task engagement. (**C**) Button push ratio as a function of the frequency ratio between the deviant and standard sounds (an example from monkey C). (**D**) Illustration of the 64-channel ECoG arrays implanted on the auditory cortex (AC) and prefrontal cortex (PFC) of monkey C (asl: arcuate sulcus lower limb; asu: arcuate sulcus upper limb; ps: principal sulcus; ls: lateral sulcus; sts: superior temporal sulcus; C: caudal; V: ventral).

We first examined neural responses to repeated standard tones and identified two distinct adaptation profiles. In the first, exemplified by an AC site, strong responses to the initial tone progressively attenuated across repetitions (Fig. 2A, top). This site showed continued attenuation through the tenth repetition, consistent with the phenomenon of *repetition suppression*, a reduction in neural response under high certainty of repeated input^72^. Time-frequency analysis revealed parallel decreases in power around 10 Hz and 2 Hz (Fig. 2A, bottom). In contrast, a second response type exhibited progressive enhancement in amplitude from the third to tenth tone (Fig. 2B, top), accompanied by rising ∼2 Hz oscillatory power aligned with the stimulus rhythm (Fig. 2B, bottom). This pattern suggests the phenomenon of *repetition enhancement*, reflecting the buildup of temporal expectations or short-term memory of the tone sequence^73^. Finally, a representative PFC site showed minimal responses to early repetitions (3–6) but strong activation during later ones (8–10), paralleled by increased ∼2 Hz power (Fig. 2C), indicating that prediction-related dynamics also emerged in frontal regions.

**Figure 2.**
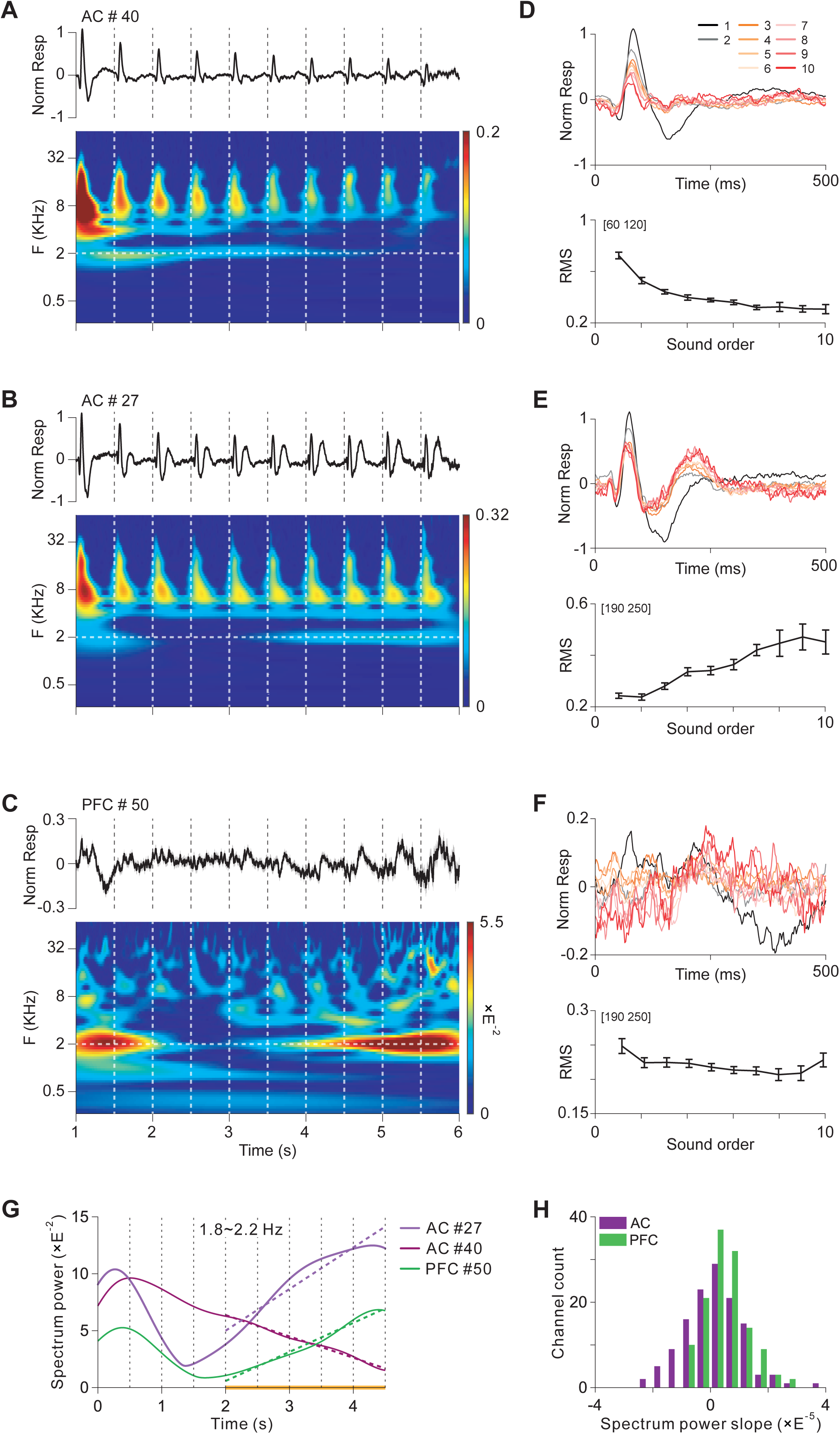
Suppression and enhancement of prediction in the AC and PFC of the monkey. (**A**) Example AC channel (#40) from monkey X showing prediction suppression. The top panel shows the normalized response averaged across trials, while the bottom panel shows the corresponding time-frequency response, where color represents spectrum power. Vertical dashed lines indicate the onset of each standard sound, and the horizontal dashed line marks the 2-Hz frequency corresponding to the 500-ms ISI. (**B-C**) Example channels (B: AC #27, C: PFC #50) from monkey X showing prediction enhancement. (**D-F**) Top: normalized responses of standard sounds from orders 1 to 10 for the three example channels. Bottom: root mean square (RMS) of the responses as a function of sound order (over the time window on the top-left). (**G**) Spectrum power in the 2-Hz frequency band (1.8**-**2.2 Hz) for the example channels, showing changes over time. Vertical black dashed lines indicate the onset of sounds. Slanted dashed lines represent linear fits from 2 to 4.5 s (corresponding to orders 5 to 9, indicated by the horizontal yellow line). (**H**) Histogram of the slopes from the linear fits in (G) for all channels in the AC and PFC for both monkeys.

To further characterize repetition-related dynamics, we aligned neural responses by presentation order. In the example AC site, evoked activity around 100 ms after tone onset progressively declined with repetition (Fig. 2D). In contrast, a second AC site (Fig. 2E) and a PFC site (Fig. 2F) showed increasing responses centered near 220 ms. These divergent profiles reveal systematic, site-specific modulation by repetition—decreasing activity in one AC site, increasing in another, and monotonic enhancement in PFC. Corresponding time–frequency maps showed parallel trends, particularly within the ∼2 Hz band (bottom rows in Figs. 2A–C).

We quantified these effects by tracking power in the 1.8–2.2 Hz band, aligned with the 2 Hz stimulation rhythm corresponding to the 500-ms ISI (Fig. 2G). Linear fits (dashed lines, Fig. 2G) across the 5^th^–9^th^ tones revealed a negative slope for the first AC site, consistent with repetition suppression, and positive slopes for the second AC and PFC sites, consistent with repetition facilitation. Across sites, 57% of AC and 76% of PFC channels showed increasing 2-Hz power over repetitions (Fig. 2H). The average slope of 2-Hz power was significantly steeper in PFC than in AC (*F*(1,254) = 14.22, *p* < 0.001, η² = 0.05; two-way ANOVA), indicating stronger low-frequency entrainment with repetition in PFC. Together, these results show that predictive facilitation dominates in prefrontal regions, reflected in the temporal buildup of low-frequency neural activity.

After characterizing local responses to standard stimuli, we next examined large-scale interactions between AC and PFC during the repetition period using Granger causality analysis. We focused on the 1–3 s window of each tone block (third–seventh tones) and averaged signals across all 64 electrodes per region (Fig. 3A). GC spectra revealed a prominent ∼2 Hz peak in both directions (AC→PFC and PFC→AC) indicating bidirectional coupling aligned with the stimulus rhythm.

**Figure 3.**
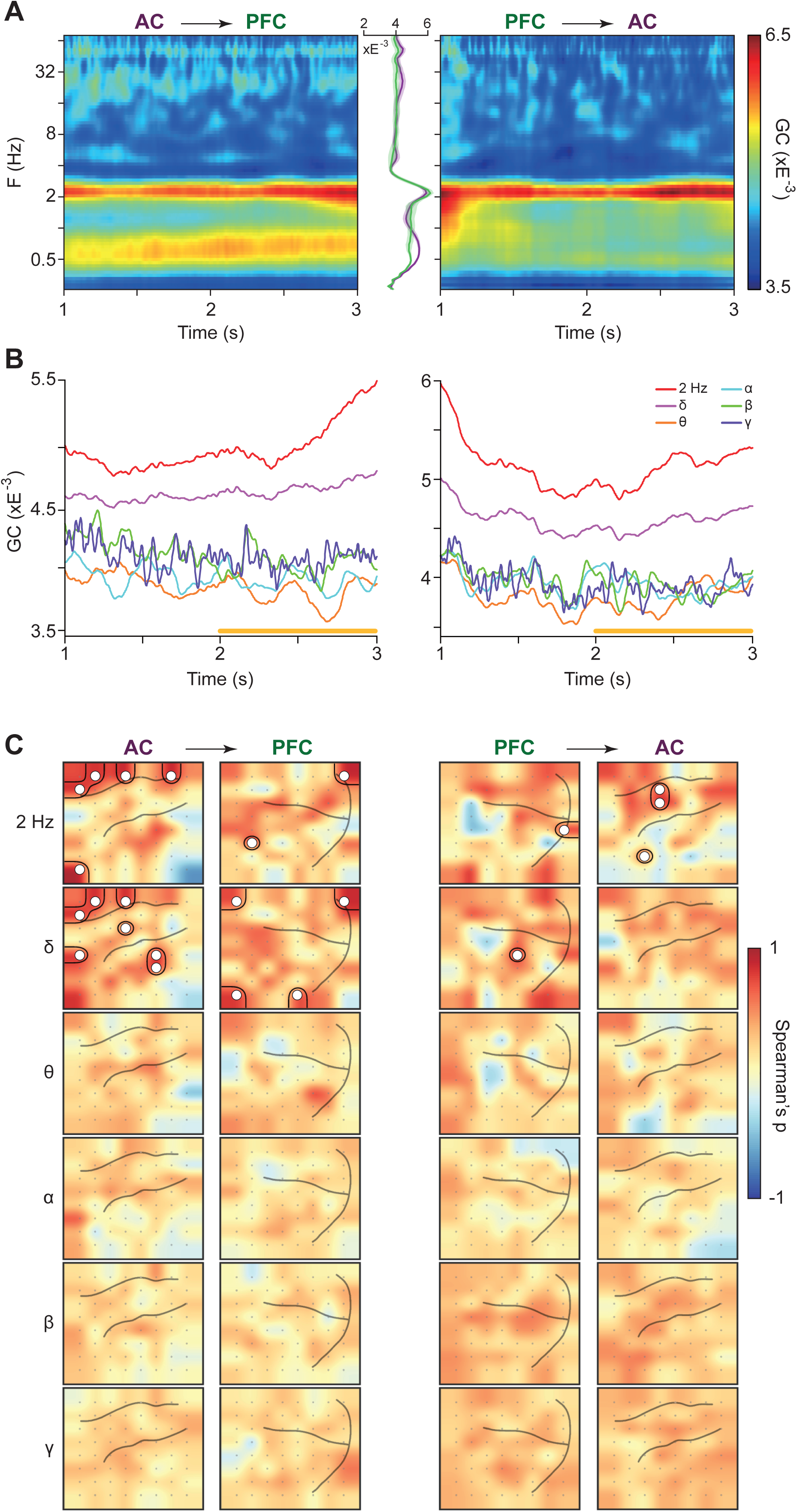
Enhancement of 2-Hz information flow between AC and PFC in prediction. (**A**) Time-frequency maps of Granger Causality (GC) spectrum, aligned to the onset of the first standard sound and averaged across all channels for monkey C. The example GC was computed using trials with a standard sound number of 8. Traces on the right and left display the GC averaged across time from 1 to 3 s, with a peak around 2 Hz. (**B**) GC for different frequency bands as a function of time, averaged across all channels for monkey C in both directions. Horizontal yellow bars indicate time window (2–3 s, orders 5–6) used for correlation calculation. (**C**) Topographic maps of GC correlation over time under each frequency band for monkey C. Channels with significant correlation are circled in black and marked with ‘o’ (Spearman’s rank correlation, *p* < 0.05, false discovery rate [FDR] corrected for channels).

To assess frequency-specific communication, we computed GC across canonical bands (2 Hz, delta: 0.6–4 Hz; theta: 4–8 Hz; alpha: 8–14 Hz; beta: 14–30 Hz; gamma: 30–70 Hz) (Fig. 3B). In both, AC→PFC (bottom-up) and PFC→AC (top-down) directions, connectivity was strongest at 2 Hz and within the delta range, with minimal coupling at higher frequencies. Temporal correlations of GC (Spearman, 2–3 s window) confirmed that significant, stable interactions emerged exclusively in the 2 Hz and delta bands (Fig. 3C). These effects were consistent across animals (Fig. 3C and Supplementary Fig. 3). Notably, when the repetition rate was increased to 2.5 Hz, GC peaks remained near 2 Hz (Supplementary Fig. 4A), suggesting that delta-band coupling reflects an intrinsic resonance of the AC–PFC circuit rather than simple entrainment to stimulus timing.

### ‘Prediction error’ mechanisms during novelty detection in AC and PFC under behavioral conditions

Next, we analyzed neuronal responses to deviant tones in both AC and PFC. An example AC site showed robust deviant responses whose magnitude scaled with the frequency difference from standards (Fig. 4A). In contrast, a representative PFC site exhibited minimal activity to standards but strong activation to deviants (Fig. 4B). To quantify these effects, we extracted two measures: (1) the first significant time, defined as the earliest time bin showing a significant deviant–standard difference in a cluster-based permutation test, and (2) the maximum *F*-value before 350 ms, indexing prediction-error strength. In the illustrated AC site, the first significant time occurred at 38 ms with a maximum *F*-value of 20.93, whereas in the PFC site, significance emerged later at 144 ms with a peak *F*-value of 14.27.

**Figure 4.**
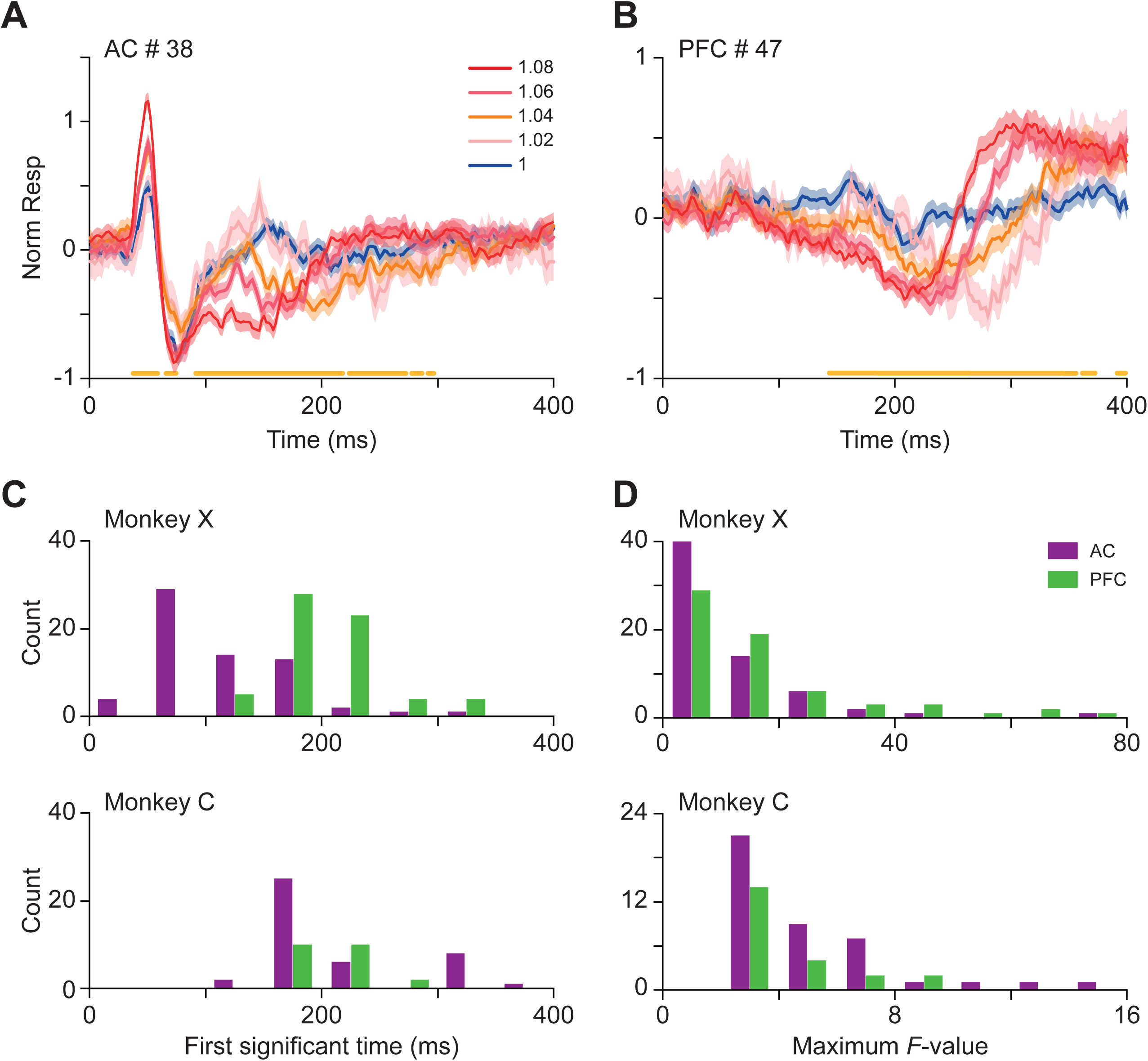
Prediction-error in the AC and PFC of the monkey. (**A**) Normalized responses for channel 38 in the AC of monkey X under five frequency ratios, aligned to the onset of the last sound and averaged across trials. Yellow bars at the bottom indicate significant differences among the 5 traces (permutation test, *p* < 0.05, FDR corrected for channels and time bins). (**B**) Same as (A) but for channel 47 in the PFC of monkey X. (**C**) Histogram of time bins when the five traces start to show significant differences for each channel. The AC showed faster predictive error than PFC (two-way ANOVA: *p* < 0.001 between areas, *p* < 0.01 between monkeys). (**D**) Histogram of *F*-values averaged from 0 to 350 ms for each channel. No significant difference was found between areas for both monkeys (Kruskal-Willis test: *p* = 0.13 for monkey X, *p* = 0.2 for monkey C).

At the population level, deviant–standard differences emerged reliably earlier in AC than in PFC for both monkeys (Fig. 4C; monkey X: *Z* = 7.67, *p* < 0.001, *r* = 0.68; monkey C: *Z* = 2.32, *p* = 0.02, *r* = 0.29; two-tailed Mann–Whitney U test, AC vs. PFC).

In contrast, maximum *F*-values did not differ significantly between regions (Fig. 4D; monkey X: *Z* = 1.48, *p* = 0.14, *r* = 0.13; monkey C: *Z* = 0.68, *p* = 0.49, *r* = 0.09).

Sorting channels by first significant time further confirmed earlier deviance detection in AC (Supplementary Figs. 5A, B). Spatial maps of maximum *F*-values revealed heterogeneous distributions within both AC and PFC, suggesting localized or tonotopic organization of prediction-error signaling (Supplementary Figs. 5C, D).

To probe functional connectivity during deviant processing, we applied Granger causality analysis. Time-frequency GC spectra (aligned to stimulus onset and averaged across channels in monkey X) revealed weak directional coupling during standards (Fig. 5A, top) but a marked GC increase for deviants (1.08 frequency ratio; Fig. 5A, middle). This enhancement emerged within 100 ms, dominated by delta/theta activity (0.6–8 Hz, AC→PFC), and shifted to theta/alpha frequencies (4–14 Hz, PFC→AC) around 200–300 ms (Fig. 5A, bottom), indicating rapid, frequency-specific engagement of the AC–PFC network during prediction-error signaling.

**Figure 5.**
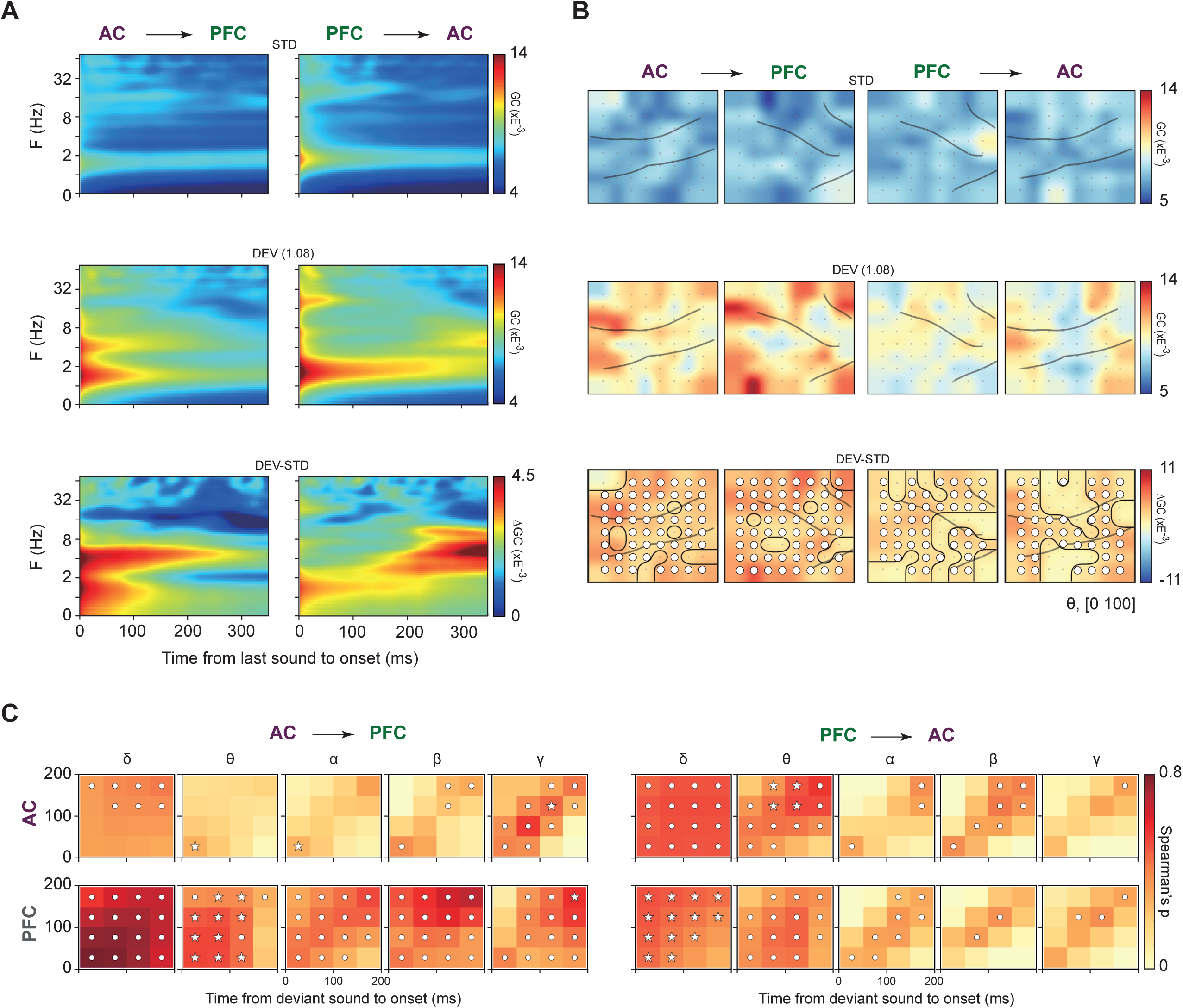
Information flow of prediction-error between the AC and PFC of the monkey. (**A**) Time-frequency maps of GC spectrum, aligned to the onset of the last sound and averaged across all channels for monkey X. Left: from AC to PFC. Right: from PFC to AC. Top: last sound as standard sound. Middle: last sound as deviant sound with a frequency ratio of 1.08. Bottom: differential GC (deviant 1.08 subtracted by standard). (**B**) Topographic GC maps for standard (top), deviant (middle), and differential (bottom) conditions, averaged in the theta band and the [0 100] ms time window. Channels with significant differences between responses to deviant and standard sounds are circled in black and marked with ’o’ (one-tailed permutation test, *p* < 0.05, FDR corrected for channels). (**C**) Averaged cross-period correlation matrices of differential topographic GC maps between pairs in frequency ratios of 1.04, 1.06, and 1.08 for monkey X. The color scale indicates Spearman’s ρ. Significant correlations for all pairs are indicated by dots for one monkey and by stars for both monkeys (Spearman’s rank correlation, *p* < 0.05, uncorrected).

Topographic GC maps corroborated these effects (Fig. 5B): bidirectional coupling was minimal for standards but rose sharply for deviants, particularly in the AC→PFC direction, with subtraction maps confirming the strongest increases along this pathway. Across frequency bands, significant information flow spanned delta to gamma ranges in both directions and in both monkeys (Supplementary Fig. 6). Temporal correlation matrices of GC values across 64 channels and frequency-ratio pairs (1.04 vs. 1.06, 1.04 vs. 1.08, 1.06 vs. 1.08) further showed robust AC→PFC interactions in the theta/alpha (4–14 Hz) and gamma (30–70 Hz) bands, while PFC→AC coupling predominated in the delta and theta (4–8 Hz) ranges (Fig. 5C). These connectivity patterns were reproduced in the second monkey (Supplementary Fig. 7), confirming dynamic, bidirectional coordination between auditory and prefrontal cortices during deviant detection.

### Active behavioral states amplify predictive and prediction-error dynamics in AC and PFC

In the previous sections, we characterized how prediction and prediction-error signals emerged during active behavioral engagement. However, to determine whether these computations required task involvement or instead reflected more general auditory processing, it was essential to examine them under passive listening as well (Fig. 1B). Therefore, we next compared neural responses during passive presentation of the auditory sequences with those recorded while monkeys performed the active novelty-detection task, allowing us to assess how behavioral engagement shaped prediction and prediction-error processing. We first examined predictive processing. In the behavioral condition (Fig. 1A), a representative PFC site displayed robust ∼2 Hz oscillatory activity between 3.5 and 5 s (Fig. 6A, left), which was markedly reduced during passive listening (Fig. 6A, right). To assess regional specificity, we compared all 64 recording sites within each area (Fig. 6B). Behavioral modulation of prediction-related activity was stronger in PFC than in AC, as shown by site-by-site scatter plots (Fig. 6C). Statistical analysis confirmed a significant behavioral effect in PFC (*F*(1,126) = 12.45, *p* < 0.001, η² = 0.05; Fig. 6C, bottom), but not in AC (*F*(1,126) = 1.43, *p* = 0.23, η² = 0.01; Fig. 6C, top). Granger causality analysis further revealed weaker AC–PFC connectivity during passive listening (Supplementary Fig. 8A) compared with active engagement (Fig. 3A). Unlike the pronounced bidirectional dynamics observed during task performance, temporal modulation of connectivity was largely absent under passive conditions (Supplementary Fig. 8B).

**Figure 6.**
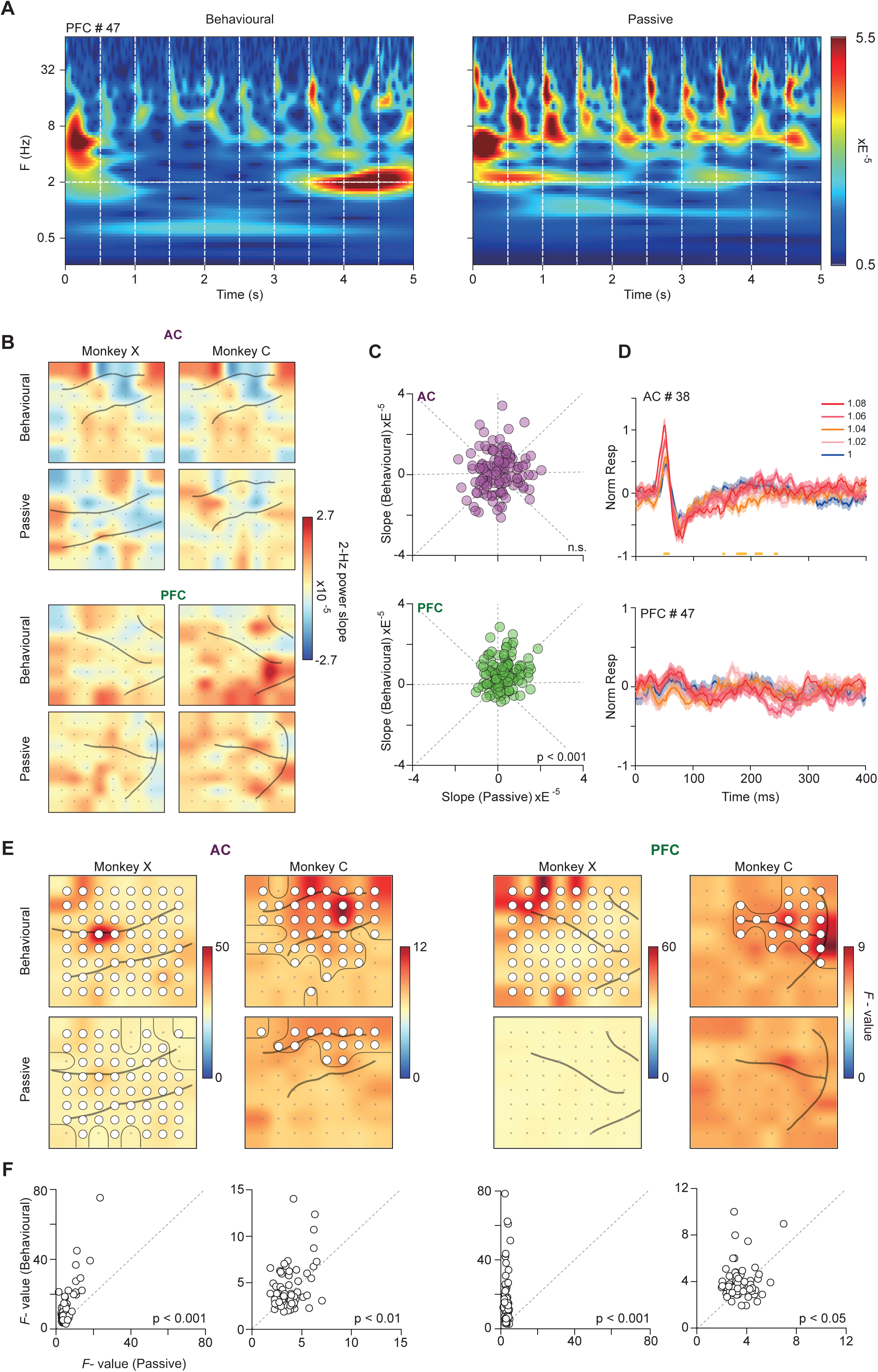
Behavioral context modulates prediction and prediction-error. (**A**) Time-frequency response of an example PFC channel during standard stimulus presentation under behavioral (left panel) and non-behavioral (right panel) contexts. (**B**) Topographic maps show the slopes of linear fits of 2-Hz power against time bins (from 5^th^ to 9^th^ standard sounds) in AC and PFC under different contexts for both monkeys. (**C**) Scatterplots comparing slopes of 2-Hz power under behavioral and non-behavioral context, with each circle representing one recording site. The results of both monkeys are plotted together. Two-way ANOVA showed a main effect of behavioral context in PFC (*p* < 0.001) but not in AC (n.s., nonsignificant). No main effect of monkey was found for both areas. (**D**) Normalized responses for example channels from monkey X (same channel displayed in Figs. 4A**-**B) under five frequency ratios, aligned to the onset of the last sound and averaged across trials. Yellow bars at the bottom indicate significant differences among the 5 traces (permutation test, *p* < 0.05, FDR corrected for channels and time bins). No significant difference was found in PFC. (**E**) Topographic maps of maximum *F*-values within 350 ms post-stimulus for two areas for both monkeys. Channels with significant difference (< 350 ms) are circled in black and marked with ‘o’. (**F**) Scatterplots comparing maximum *F*-values under behavioral and non-behavioral contexts, with areas and monkeys aligned with (E). Two-tailed paired *t*-test showed significant increase of *F*-values under behavioral context for both areas and monkeys—AC of monkey X: *t*(63) = 5.47, *p* < 0.001, *d* = 0.68; AC of Monkey C: *t*(63) = 2.51, *p* < 0.05, *d* = 0.31; PFC of monkey X: *t*(63) = 6.68, *p* < 0.001, *d* = 0.84; PFC of monkey C: *t*(63) = 2.1, *p* < 0.05, *d* = 0.26.

We next examined prediction-error signaling. Example sites in both AC and PFC (Fig. 6D, top and bottom, respectively) showed markedly reduced differentiation between deviant and standard tones during passive listening, compared with the clear response separation observed during active task performance (*cf.*, Figs. 4A, B). Across both regions and animals, the tonotopic distribution of *F*-values was attenuated in the passive condition (Fig. 6E). Site-wise comparisons confirmed significantly stronger prediction-error responses during active behavior (*p* < 0.05 for all comparisons; two-tailed paired t-test; Fig. 6F). Granger causality analysis nonetheless revealed residual AC→PFC information flow in the theta and alpha bands, indicating that basic feedforward signaling persisted even without behavioral engagement (Supplementary Fig. 9).

## Discussion

In this study, we examined how the auditory and prefrontal cortices cooperate to predict and detect sensory novelty in the auditory cortex and prefrontal cortex of behaving macaques. Repetition of standard tones produced two distinct response profiles: while some AC sites showed repetition suppression, most AC and PFC sites exhibited repetition enhancement, dominated by low-frequency (∼2 Hz) oscillatory activity (Fig. 2). Linear fits of delta-band power confirmed stronger facilitation in PFC than in AC (Fig. 2H). Granger causality analyses revealed robust bidirectional coupling between the two cortical regions, peaking near 2 Hz within the delta band, consistent with coordinated predictive signaling (Fig. 3; Supplementary Figs. 3–4). Deviant tones elicited early responses in AC and later activation in PFC (Figs. 4A–C), with comparable prediction error magnitudes across the cortical network (Fig. 4D). Deviance processing further increased AC→PFC information flow in the theta-alpha bands and induced frequency-specific cross-cortical coupling, gamma from AC to PFC and delta–theta from PFC to AC (Fig. 5C; Supplementary Figs. 6–7). Importantly, both predictive and error related responses were amplified during active engagement (Fig. 7), showing enhanced repetition enhancement, cross-cortical connectivity, and deviance sensitivity compared with passive listening (Fig. 6; Supplementary Figs. 8–9). Together, these results support a distributed, state-dependent model of predictive coding in which the AC encodes early sensory regularities and violations, whereas the PFC integrates behavioral context to shape top-down modulation (Fig. 7).

**Figure 7.**
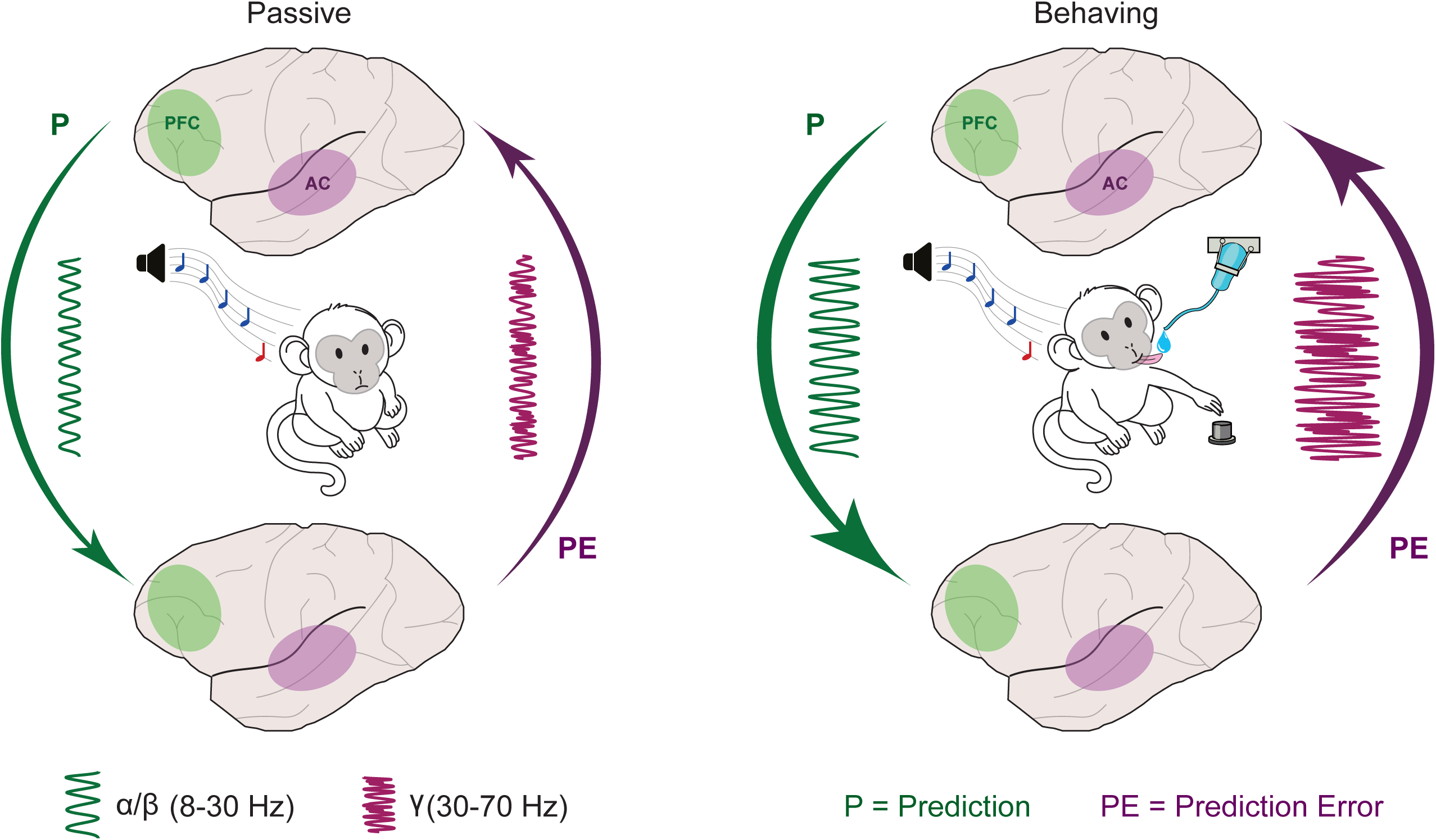
State-dependent predictive coding between auditory and prefrontal cortex during novelty detection. Schematic summary of the main findings. In both passive (left) and behaving (right) conditions, auditory cortex (AC, purple) and prefrontal cortex (PFC, green) form a recurrent predictive network. Top-down predictions (P) from PFC to AC are carried mainly by low-frequency activity, including delta (∼2 Hz) and α/β (8–30 Hz) oscillations. Bottom-up prediction errors (PE) from AC to PFC are expressed in higher-frequency θ (4–8 Hz) and γ (30–70 Hz) bands. In the passive state, prediction signals and AC–PFC coupling are weak, and repetition suppression dominates in AC. During active novelty detection, behavioral engagement amplifies prediction and prediction-error signals, enhances delta-band coupling, particularly strengthens θ/γ-band AC→PFC connectivity, and increases repetition enhancement in both regions. The circuit highlights a hierarchical, state-dependent predictive coding mechanism in which AC encodes sensory regularities and violations, while PFC integrates behavioral context to modulate auditory processing.

### Dual modes of prediction in the AC–PFC circuit

Repetition suppression and repetition enhancement represent complementary forms of predictive plasticity within the auditory–prefrontal cortices network. In the AC, repetition suppression (observed in ∼43% of electrodes) appeared as a gradual attenuation of evoked responses to repeated tones (Figs. 2A, D), consistent with predictive coding models in which redundant input reduces prediction-error signaling in primary auditory and subcortical nuclei^4,5,72,74^. The remaining AC sites, however, exhibited pronounced repetition enhancement, with evoked amplitudes and phase-locked power at the stimulation rhythm (∼2 Hz) increasing monotonically across repetitions (Figs. 2B, E). Similar facilitation reported in marmoset and mouse AC^75,76^ may represent a transient “prediction template” that sharpens temporal expectancy through delta-phase alignment^77^. The PFC showed the same dual motifs but was strongly biased toward enhancement: 76% of electrodes displayed progressive facilitation, typically with a later buildup than in AC (Figs. 2C, F–H). This predominance accords with evidence that medial PFC broadcasts precision-weighted predictions to increase auditory gain under high behavioral relevance^50,78^ and sustains cross-modal predictive templates during working memory^78^.

Interestingly, these predictive effects were strongly modulated by behavioral state. During passive listening, repetition enhancement strength declined sharply in PFC (33% reduction in the average 2-Hz power slope relative to the active condition) and more moderately in AC (22%), accompanied by weakened low-frequency PFC–AC coupling (Supplementary Fig. 8). This pattern mirrors evidence that task engagement amplifies frontal contributions to sensory processing^79,80^, suggesting that PFC-driven repetition enhancement refreshes predictive templates during active behavior. In contrast, repetition suppression predominated under passive conditions, reflecting efficient suppression of redundant input and reduced resource allocation. This state dependence highlights a functional dissociation repetition suppression within the AC–PFC circuit: repetition suppression optimizes coding efficiency for predictable repetitive stimuli, whereas repetition enhancement enhances the representation of behaviorally relevant information through active cortical buffering.

Converging human magnetoencephalogram (MEG) and functional magnetic resonance (fMRI) work indicates that such dual repetition effects are a general feature of auditory memory-trace and prediction formation. Using a roving-standard paradigm, researchers^81^ have showed that repetition suppression of the N1m response is accompanied by a progressive repetition enhancement of a later sustained field, and that both components contribute to the build-up of the repetition positivity^73^ and the echoic memory trace. Another event-related fMRI study^82^ further demonstrated that deviant sounds preceded by a longer sequence of standard tones evoke stronger blood oxygen level-dependent responses in bilateral auditory cortex, and that responses to the standards themselves show a mixture of suppression and enhancement depending on whether they predict an upcoming deviant. These findings support the view that repetition suppression primarily reflects reduced prediction error for highly expected input, whereas repetition enhancement indexes the strengthening of short-term auditory predictions and sensory memory representations in auditory cortex.

Our macaque ECoG data extend this perspective in several important ways. First, we show that repetition suppression and enhancement coexist not only within AC but also in PFC, and that their relative expression is strongly state-dependent, with enhancement dominating under active task engagement. Second, the slow build-up of low-frequency activity we observe in PFC—and in a subset of AC sites—resembles the sustained-field repetition enhancement and deviant-history effects described in human MEG and fMRI, suggesting a common mechanism whereby frontal feedback and local delta-band entrainment maintain short-term auditory predictions during regular stimulation^81,82^. Third, by combining these repetition effects with directed connectivity analyses, we show that they are embedded in a structured AC–PFC network, in which PFC feedback preferentially amplifies enhancement-related dynamics under behavioral demands. This cross-species convergence supports the notion that dual repetition mechanisms—suppression and enhancement—are a core implementation of predictive coding in the auditory system, jointly supporting efficient encoding of predictable input and robust maintenance of context-dependent expectations.

The coexistence of repetition suppression and repetition enhancement within the same regions supports a flexible predictive coding framework. repetition suppression in AC reflects prediction error minimization at early sensory stages, whereas repetition enhancement in PFC indexes precision-weighted model updating^72^. Delta-phase–driven repetition enhancement in AC (Fig. 2) and task-dependent facilitation in PFC (Fig. 6) suggest that temporal coding and fronto-sensory interactions jointly refine predictive accuracy. Together, these mechanisms show how the brain balances efficiency through suppression and precision through enhancement, adapting dynamically to behavioral demands via repetition suppression, repetition enhancement, and AC–PFC coordination. Future work should examine how these processes generalize to naturalistic contexts and to clinical conditions with impaired predictive processing.

The GC patterns observed during standard stimulus repetitions revealed a structured interplay between AC and PFC that supports predictive maintenance. Bidirectional coupling peaked near 2 Hz (Figs. 3A, B), matching the stimulation rhythm and phase-locked delta oscillations in both regions (Figs. 2G, H). This delta-band coherence indicates that AC and PFC synchronize to encode temporal regularities, with reciprocal AC→PFC and PFC→AC signaling reinforcing the stability of predictive templates. The dominance of delta frequencies in these exchanges (Figs. 3B, C) aligns with theories positing that low-frequency oscillations coordinate hierarchical predictions^51,77,83^, whereby AC conveys sensory-derived temporal structure to PFC, which in turn modulates auditory gain to emphasize behaviorally relevant patterns^50^.

Task engagement strongly modulated the strength and direction of functional coupling between AC and PFC. During active performance, top-down PFC→AC influence increased markedly compared with passive listening (Supplementary Fig. 8), mirroring the higher prevalence of repetition enhancement in PFC under behaviorally relevant conditions. This asymmetry suggests that prefrontal feedback amplifies sensory predictions when sustained attention to auditory regularities is required. In contrast, passive listening reduced PFC→AC drive, shifting the balance toward bottom-up AC-driven interactions. Such flexibility accords with predictive-coding frameworks in which precision weighting—shaped by behavioral relevance—dynamically regulates the balance between top-down predictions and bottom-up sensory evidence^72,84^.

The persistence of delta-band coupling across repetition rates (Supplementary Fig. 4) further underscores its role in maintaining temporal predictions. Unlike higher-frequency interactions linked to local processing or prediction-error signaling^85^, delta oscillations appear to provide a scaffold for cross-cortical predictive maintenance. This interpretation aligns with evidence that delta-phase coding in AC sharpens temporal expectancy^77,86^, while PFC leverages these rhythms to sustain working-memory representations^87^. Together, AC–PFC delta synchrony during repetition emerges as a dynamic *prediction loop* that stabilizes auditory regularities and optimizes processing for anticipated inputs.

### Prediction-error hierarchy and AC–PFC communication

Our findings outline a hierarchical architecture for prediction-error processing in the auditory system, defined by dynamic interactions between AC and PFC. Deviant tones evoked significant responses in both regions, yet their temporal dynamics diverged: deviations from standard responses emerged ∼30–100 ms earlier in AC than in PFC (Figs. 4A–C; Supplementary Figs. 5A, B). This temporal lead aligns with evidence that prediction errors originate in lower-order auditory areas before propagating to higher-order regions, consistent with hierarchical predictive processing^88,89^. Despite this early onset, the overall magnitude of prediction-error activity—indexed by peak *F*-values—did not differ significantly between AC and PFC (Fig. 4D). These results suggest that while AC rapidly flags sensory deviations, PFC contributes to their amplification and integration, linking error detection to higher cognitive functions such as attention and decision-making^90^.

To clarify the directional flow of information during deviant processing, we examined connectivity dynamics across frequencies. Deviant stimuli triggered a rapid increase in AC→PFC coupling within the delta and theta bands, emerging within 100 ms of stimulus onset (Figs. 5A-C). This pattern reflects feedforward transmission of prediction-error signals from sensory to frontal regions. Shortly thereafter, enhanced PFC→AC coupling in the same frequency range indicated feedback processes likely involved in updating predictive models^88^. The strengthening of prefrontal feedback following deviant detection suggests a rapid recalibration of predictive templates, consistent with evidence linking frontal activity to expectation updating after surprising auditory events^88^.

### Behavioral engagement emerged as a key determinant of prediction-error dynamics

During active task performance, both AC and PFC exhibited stronger deviant responses and broader spatial differentiation of prediction-error activity than during passive listening (Figs. 6C–F). Connectivity analyses showed enhanced delta- and theta-band coupling under behavioral conditions, particularly in the AC→PFC direction, indicating greater feedforward transmission of errors when auditory processing was task-relevant (Supplementary Fig. 9). These results converge with conceptual models in which prediction-error signaling is weighted by attention and task demands^91^, highlighting behavioral relevance as a key factor shaping both the magnitude and direction of information flow across cortical networks (Fig. 7).

Together, the data support a bidirectional model of auditory predictive processing: AC rapidly detects sensory deviations and conveys them to PFC, which integrates contextual information before sending refined predictions back to sensory cortex. This recurrent exchange promotes adaptive perception and exemplifies the hierarchical reciprocity central to predictive coding theories.

## STAR Methods

### Key Resources Table

| RESOURCE | SOURCE | IDENTIFIER |
| --- | --- | --- |
| Rhesus macaque<br>(*Macaca mulatta*) | This study | N/A |
| 64-channel ECoG array | NeuroNexus | E642500-50-200 |
| RZ5 amplifier | Tucker-Davis Technologies | RZ5 |
| RZ6 processor | Tucker-Davis Technologies | RZ6 |
| MATLAB R2021a | MathWorks | RRID:SCR_001622 |
| FieldTrip toolbox | Oostenveld *et al.*, 2011 | RRID:SCR_004849 |
| BayesFactor toolbox | Rouder *et al.*, 2009 | Zenodo:<br>[10.5281/zenodo.13744717](https://zenodo.org/record/13744717) |
| Custom MATLAB<br>analysis code | This study | Github:<br>[TOMORI233/Code-for-monkey-AC-PFC](https://github.com/TOMORI233/Code-for-monkey-AC-PFC) |
| Raw and processed<br>ECoG data | This study | Zenodo:<br>[10.5281/zenodo.18017973](https://zenodo.org/record/18017973) |

### Resource Availability

#### Lead contact

Further information and requests for resources should be directed to and will be fulfilled by the lead contact, Xiongjie Yu.

#### Materials availability

This study did not generate new unique reagents.

#### Data and code availability

- All raw and processed ECoG data have been deposited at Zenodo (DOI: 10.5281/zenodo.18017973) and were deposited prior to submission.
- Custom MATLAB codes are available at GitHub (https://github.com/TOMORI233/Code-for-monkey-AC-PFC.git).
- Any additional information required to reanalyze the data reported in this paper is available from the lead contact upon reasonable request.

#### Experimental Model and Subject Details

We conducted experiments in two adult male rhesus monkeys (Macaca mulatta; subjects C and X), aged 7 years and weighing between 5.5 and 7 kg. All experimental protocols strictly complied with the Guide for the Care and Use of Laboratory Animals (Directives 86/609/EEC, 2003/65/EC, and 2010/63/EU) and the Regulations for the Administration of Affairs Concerning Experimental Animals of the People’s Republic of China (GB 14925-2010), and were approved by the Bioethics Committee of Zhejiang University (ZJU20200148). Throughout the study, researchers and animal care staff closely monitored both monkeys daily to ensure their health and well-being. To enhance their quality of life, toys containing food items that the monkeys enjoyed were routinely introduced into their 0.74 m^3^ home cage to promote exploratory behavior. Regular assessments for physical and physiological well-being of the animals were carried out by veterinarians, registered veterinary technicians and researchers.

Each monkey was implanted with two 64-channel subdural electrode arrays (E64-2500-50-200, NeuroNexus Technologies, USA) positioned over the left auditory cortex (AC) and prefrontal cortex (PFC). The surgical and ECoG recording procedures followed those described in our previous report^92^.

Aseptic techniques were employed throughout surgery. Monkeys were premedicated with ketamine (50 mg/kg) and medetomidine (0.03 mg/kg), then intubated and maintained under anesthesia with artificial ventilation to stabilize body temperature at 37°C using an electric heating mat. Vital signs (oxygen saturation, heart rate, and end-tidal CO_2_) were continuously monitored to adjust anesthesia depth as required. Each subject’s head was stabilized in a stereotaxic frame (Narishige, Japan). After local infiltration with lidocaine, the scalp and overlying muscles were retracted, and a titanium headpost (Gray Matter Research, USA) was secured to the skull with bone screws and resin. Craniotomy and durotomy were performed under a surgical microscope (Carl Zeiss, Germany). Following implantation, the dura mater, bone flap, and skin were sutured, and the exposed area was covered with resin. Postoperative care included ketoprofen administration for three days and antibiotic treatment for one week to prevent infection and ensure recovery.

The ECoG electrodes had a diameter of 0.2 mm and gold recording sites embedded in a 2 × 2 cm polyimide substrate. Electrode impedances measured at 1 kHz ranged from 20 to 50 kΩ, with an inter-electrode spacing of 2.5 mm. Preoperative MRI scans were used to determine the target coordinates and craniotomy size for each subject. A gold reference electrode was placed near the ECoG array within the subdural space and oriented toward the dura mater. Lead wires from both the ECoG and reference electrodes were connected to micro-connectors (ZIF-Clip 64, Tucker-Davis Technologies, USA), housed in a titanium chamber fixed to the skull with dental resin. Neural signals were amplified and band-pass filtered between 0.5 and 300 Hz using a differential amplifier (RZ5, Tucker-Davis Technologies).

## Methods Details

### Auditory stimulus generation

All recordings were performed in a sound-attenuated chamber. Monkeys were seated in a primate chair with the head fixed, the right ear oriented toward a free-field loudspeaker (LS50, KEF, UK), and one hand resting on a response button. Acoustic stimuli were digitally generated using a computer-controlled auditory workstation (RZ6, Tucker-Davis Technologies) at a 100 kHz sampling rate and delivered through the contralateral (right) speaker. Sound pressure levels were calibrated using a ¼-inch condenser microphone (Brüel & Kjær 4954, Denmark) connected to a PHOTON/RT analyzer (Brüel & Kjær, Denmark).

The stimuli consisted of complex tones composed of seven harmonics, defined by the following equation:

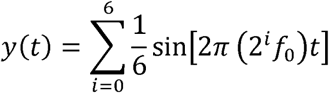

where *t* represents time and *f*_0_ is the base frequency of the tone. The standard sound had a base frequency of 245 Hz. For deviant tones, the base frequency was altered relative to the standard according to the ratio λ, such that *f*_dev_ = *f*_std_ x *λ*, with *λ* randomly selected from 1.02, 1.04, 1.06, or 1.08. Both standard and deviant tones had a duration of 100 ms and were presented at 60 dB SPL.

### Behavioral paradigm

Monkeys were trained to perform a novelty-detection task based on an auditory oddball paradigm (Fig. 1)^41,42,93^. Each trial began when the monkey pressed a button to initiate the sequence. Two seconds after the press, a series of 8–11 complex tones were presented at a rate of 2 Hz (500-ms inter-stimulus interval (ISI), defined as the time between sound onsets). A 5-s inter-trial interval was imposed, during which button presses were invalid.

Within each sequence, the initial tones were standards with identical frequencies, while the final tone served as the target. In deviant trials, the target tone differed in frequency from the preceding sounds, whereas in control trials it was identical. The monkey was required to press the button within 600 ms after deviant onset to receive a water reward. This response window prevented random button presses at the end of each block, since the minimum reaction time exceeded 200 ms.

Deviant trials were defined by the frequency ratio (λ) between deviant and standard tones, with λ = 1.02, 1.04, 1.06, or 1.08. Control trials corresponded to λ = 1. Each condition was presented randomly and repeated 40–50 times per recording session. The number of standards preceding the target varied randomly between 7 and 10 to prevent anticipatory responses based on sequence length.

After each behavioral session, a non-behavioral session was conducted in which a baffle was placed between the button and the monkey’s hands to block any motor responses. Additionally, an extra behavioral session was recorded using a shorter ISI of 400 ms for both monkeys.

Behavioral performance was evaluated according to two criteria. First, Correct Hit (Deviant Identification): the monkey had to correctly identify the deviant sound with at least 80 % accuracy (button-press ratio > 0.8) for the largest frequency difference condition. Second, Correct Rejection (Absence of Deviant): in control trials, the monkey had to correctly withhold responses, achieving at least 90 % accuracy (button-press ratio < 0.1). Only recording sessions meeting both criteria were included in the analyses. In total, five behavioral and four non-behavioral sessions were recorded per monkey.

### ECoG preprocessing

ECoG data were downsampled to 500 Hz and band-pass filtered between 0.1 and 200 Hz, followed by a 50 Hz notch filter to remove line noise. The filtered signals were segmented from 3 s before to 7 s after the onset of the first sound in each trial. To reduce noise and artifact contamination, we performed independent component analysis (ICA) using the *runica* function from the *FieldTrip* toolbox^94^. Components were visually inspected and retained or rejected based on their spatial topography and variance.

After ICA, the data were re-referenced in three consecutive steps. First, we applied common average referencing (CAR) by subtracting the mean signal across all channels within each array. This step minimized inter-array differences related to electrode impedance and array-specific properties. Second, we applied orthogonal source derivation by subtracting the average signal from the four nearest neighboring electrodes (two horizontal and two vertical). This local re-referencing reduced residual common-reference and volume-conduction effects, thereby enhancing spatial specificity. Finally, the signals were normalized by their temporal standard deviation to ensure equal weighting across electrodes and trials. The re-referenced data were then averaged across recording sessions for subsequent response comparison and time–frequency analysis.

For Granger causality (GC) analysis, we applied an additional preprocessing step to the re-referenced data. The average event-related potential (ERP) was subtracted from individual trials to remove phase-locked evoked components associated with stimulus onsets and offsets. This procedure isolated non–phase-locked, internally generated activity, such as induced oscillations, which better reflect functional interactions between cortical regions.

### Time-frequency decomposition

Time-frequency analysis was performed on both re-referenced and ERP-subtracted datasets using the MATLAB *cwt* function. Morlet wavelets with a time–bandwidth product of 5 (default parameters) were used as the analytic basis functions.

For prediction analyses, the time-frequency response (TFR) during standard sound presentations was computed using the integrated ERP data. ERPs corresponding to trials with different numbers of standard sounds were calculated separately and then combined using a weighted average. To minimize edge artifacts introduced by the cone of influence (COI) in the wavelet transform, data segments were zero-padded to 10 s on both sides before transformation and subsequently trimmed to their original duration.

For Granger causality analysis, TFRs were computed at the single-trial level. Specifically, in prediction GC analysis, only the standard sound period of each trial was included, from the onset of the first tone up to the onset of the deviant tone (or one ISI after the last standard tone in control trials). For prediction-error GC analysis, TFRs were computed using the 1 s period following the onset of the final deviant or standard sound.

### Repetition suppression and enhancement analysis

Prediction-related suppression and enhancement were quantified by averaging spectral power within a frequency band centered around the stimulus repetition rate (± 0.2 Hz relative to the repeat rate). This corresponded to 1.8–2.2 Hz for the 500-ms ISI and 2.3–2.7 Hz for the 400-ms ISI (Fig. 2G; Supplementary Fig. 2C). To estimate temporal changes in predictive activity, the mean spectral power at the repeat rate was fitted across the sequence of standard sounds, specifically from the 4^th^ to the 9^th^ complex tone (Fig. 2G; Supplementary Fig. 2C). The resulting slopes of these linear fits were compared between cortical regions (AC vs. PFC; Fig. 2H; Supplementary Fig. 2D), ISI conditions (400 vs. 500 ms; Supplementary Figs. 2E–F), and behavioral contexts (behavioral vs. passive; Figs. 6B–C).

### Nonparametric Granger causality analysis

To assess the directionality and frequency specificity of interareal communication between the AC and PFC, we applied a nonparametric Granger causality (GC) approach. This method quantifies directed functional interactions by testing whether past activity in one signal improves the prediction of future activity in another. Nonparametric GC estimates were obtained by factorizing the cross-spectral density (CSD) matrix, which was computed from wavelet-transformed signals for all AC–PFC electrode pairs.

For prediction-related GC analyses, all trials without button presses during the standard-sound period were included. Data segments spanned from the onset of the first sound to the onset of the deviant sound, or to one inter-stimulus interval after the final standard tone in control trials. Trials containing different numbers of standard tones (7–10) were analyzed separately. For prediction-error GC analyses, data were extracted from non-behavioral sessions and from successful behavioral trials, covering the interval from the onset of the final sound to 1 s post-stimulus.

GC values were computed bidirectionally (AC→PFC and PFC→AC) for all electrode pairs across 72 frequency bins (0.6–70 Hz) and time bins with a 2-ms step size. Baseline GC values, estimated from a pre-stimulus window (-3 to -2.5 s relative to the onset of the first sound), were subtracted from subsequent results. Frequencies were grouped into five canonical bands: δ (0.6–4 Hz), θ (4–8 Hz), α (8–14 Hz), β (14–30 Hz), and γ (30–70 Hz), and GC values were averaged within each band.

To estimate regional-level directional information flow, GC maps were generated by averaging GC values between each electrode in one cortical area (AC or PFC) and all electrodes in the other area. This procedure was applied for both directions (AC→PFC and PFC→AC), across frequency ratios (1, 1.02, 1.04, 1.06, and 1.08), frequency bands, and overlapping temporal windows (100-ms windows with 50-ms overlap).

For prediction-related analyses, temporal modulation of information flow was quantified by computing Spearman’s rank correlations of GC values across consecutive time windows corresponding to sound orders 5–6 (2–2.5 s for the 500-ms ISI condition) for each electrode and standard-tone condition. Correlation coefficients were averaged across conditions with 7–10 standard tones to generate topographic maps for each frequency band and direction. Electrodes exhibiting significant correlations (*p* < 0.05, false discovery rate–corrected across electrodes) were marked in the corresponding figures (Fig. 3C; Supplementary Figs. 3–4).

For prediction-error analyses, differential GC maps were computed by subtracting the GC map for the standard condition (frequency ratio = 1) from that of each deviant condition. Spatial consistency of differential GC patterns across deviant magnitudes was assessed by calculating Spearman’s rank correlations between differential GC maps for each pair of deviant conditions (1.04, 1.06, and 1.08) within each frequency band, separately for each monkey and behavioral context (Fig. 5C; Supplementary Figs. 7 and 9). The smallest deviant condition (1.02) was excluded due to an insufficient number of successful trials. Cross-condition similarity was considered reliable when correlations reached *p* < 0.05 (uncorrected) across all deviant-condition pairs in both animals.

### Quantification and Statistical Analysis

All statistical analyses were performed using custom MATLAB scripts (R2021a, MathWorks) in combination with the *FieldTrip* toolbox and the *BayesFactor* toolbox^95^. Unless otherwise stated, statistical significance was assessed at *p* < 0.05.

### Definition of sample size and data inclusion

Sample size (*n*) refers to the number of recording sites (electrodes) unless otherwise specified. Electrodes were treated as independent samples, consistent with prior ECoG studies; results were verified to be consistent at the session and subject levels. Analyses were performed separately for each monkey and then combined at the population level. Only recording sessions that met predefined behavioral performance criteria (see Behavioral paradigm) were included. No statistical methods were used to predetermine sample sizes; sample sizes were similar to those used in previous non-human primate ECoG studies.

### Analysis of prediction-related neural dynamics

To quantify repetition-related suppression and enhancement, spectral power within a narrow band centered on the stimulus repetition rate (± 0.2 Hz) was averaged for each electrode and fitted across successive standard sounds using linear regression. Slopes of these fits were compared across cortical regions (AC vs. PFC), behavioral contexts (behavioral vs. non-behavioral), stimulus conditions, and subjects using two-way ANOVA. Effect sizes were reported as partial eta squared (η^2^), with values of 0.01, 0.06, and 0.14 indicating small, medium, and large effects, respectively.

### Prediction-error latency and magnitude

Prediction-error latency was defined as the earliest time point at which neural responses differed significantly across frequency-ratio conditions, as determined by cluster-based permutation tests. Prediction-error magnitude was quantified as the maximum *F*-value within the first 350 ms following stimulus onset. Latencies and magnitudes were compared between cortical regions using two-tailed Mann–Whitney *U* tests, with effect sizes reported as rank-biserial correlation (*r*).

### Cluster-based permutation testing

Differences in neural responses across frequency-ratio conditions were evaluated using nonparametric cluster-based permutation tests implemented in *FieldTrip*. For each electrode and time point, an *F*-statistic was computed across conditions. Samples exceeding an uncorrected threshold of *p* < 0.05 were grouped into spatiotemporal clusters based on electrode adjacency. Cluster-level statistics were defined as the sum of *F*-values within each cluster and compared against a null distribution generated from 1,000 random permutations. Clusters were considered significant if their statistics exceeded the 95% confidence interval of the null distribution, controlling the family-wise error rate.

### Granger causality analyses

For GC analyses, baseline-corrected GC values were averaged within predefined frequency bands (δ: 0.6–4 Hz; θ: 4–8 Hz; α: 8–14 Hz; β: 14–30 Hz; γ: 30–70 Hz).

Temporal modulation of GC during prediction was assessed by computing Spearman’s rank correlations of GC values across consecutive time windows for each electrode.

Resulting correlation coefficients were tested against zero, and false discovery rate (FDR) correction was applied across electrodes.

For prediction-error analyses, differential GC maps were obtained by subtracting GC values for standard conditions from those for deviant conditions. Spatial consistency of differential GC patterns across deviant magnitudes was quantified using Spearman’s rank correlations between GC maps from different deviant conditions.

### Permutation tests for GC contrasts

To assess condition-dependent differences in GC strength, one-tailed permutation tests were performed by randomly shuffling condition labels across trials (1,000 permutations) to generate null distributions of GC differences. Electrodes were considered significant when *p* < 0.05 across all tested deviant conditions.

### Bayesian statistical analyses

Bayesian paired or independent-sample *t* tests with Jeffreys–Zellner–Siow priors^96^ were used to quantify evidence for or against the null hypothesis in pairwise comparisons. For multi-factorial designs, Bayesian ANOVA with default Cauchy priors^97^ was applied. Bayes factors (BF_10_) greater than 1 were interpreted as evidence for the alternative hypothesis, whereas BF_10_ values less than 1 indicated evidence in favor of the null.

## Supplementary Figure legend

**Supplementary Figure 1.** ECoG array implantation. (**A**) ECoG array implantation (left: before implantation, middle: after implantation) in the PFC (top) and AC (bottom) of monkey X. Right schematic shows the relative position of the electrode sites and sulcus (asl: arcuate sulcus lower limb; asu: arcuate sulcus upper limb; ps: principal sulcus; ls: lateral sulcus; sts: superior temporal sulcus; C: caudal; D: dorsal). (**B**) Same as (A) for monkey C.

**Supplementary Figure 2.** Suppression and enhancement of prediction in the AC and PFC of the monkey under the 400-ms ISI condition. (**A-B**) Example channels from monkey X showing prediction enhancement under the 400-ms ISI condition. Vertical dashed lines indicate the onset of each standard sound, and the horizontal dashed line represents the 2.5-Hz frequency corresponding to the 400-ms ISI. (**C**) Spectrum power in the 2.5-Hz frequency band (2.3–2.7 Hz) for the example channels, showing changes over time. Vertical orange dashed lines indicate the onset of sounds. Slanted dashed lines represent linear fits from 1.6 to 3.6 s (corresponding to orders 5 to 9, indicated by the horizontal yellow line). (**D**) Histogram of the slopes from the linear fits in (C) for all channels in the AC and PFC for both monkeys. No significant differences were observed between areas and between monkeys for both prediction suppression and enhancement (two-way ANOVA: *p* = 0.08 between areas, *p* = 0.12 between monkeys for suppression; *p* = 0.44 between areas, *p* = 0.06 between monkeys for enhancement). (**E**) Scatter plot comparing the slopes between the 500-ms and 400-ms ISI conditions. No significant correlation was observed between the two conditions for both areas (*p* > 0.1, Pearson’s correlation). (**F**) Topographic comparison of slopes between the 500-ms and 400-ms ISI conditions. No significant correlation was observed between the two conditions in both areas for both monkeys (*p* > 0.1, Spearman’s rank correlation).

**Supplementary Figure 3.** Enhancement of information flow between AC and PFC in prediction in monkey. **X.** Topographic maps of GC correlation over time under each frequency band for monkey X. Channels with significant correlation are circled in black and marked with ‘o’ (Spearman’s correlation, *p* < 0.05, false discovery rate [FDR] corrected).

**Supplementary Figure 4.** 2-Hz as a characteristic frequency of prediction. (**A**) Time-frequency maps of GC spectrum, aligned to the onset of the first standard sound and averaged across all channels for monkey X under the 400-ms ISI condition. Traces on the right and left display the GC averaged across time from 0.8 to 2.4 s, with a peak around 2 Hz. White dashed and red solid vertical lines indicate 2.5 Hz corresponding to the 400-ms ISI. (**B**) GC for different frequency bands as a function of time, averaged across all channels for monkey X. Horizontal yellow bars indicate time window (1.6–2.4 s, orders 5-6) used for correlation calculation. (**C**) Topographic maps of GC correlation over time under each frequency band for monkey C. Channels with significant correlation are circled in black and marked with ‘o’ (Spearman’s rank correlation, *p* < 0.05, false discovery rate [FDR] corrected).

**Supplementary Figure 5.** Statistical results of prediction-error for all channels in both monkeys. (**A**) *F*-values of response comparisons among five frequency ratios for channels in the AC (top) and PFC (bottom) of monkey X, aligned to the onset of the last sound. Colors indicate significant time bins (permutation test, *p* < 0.05, FDR corrected for channels and time bins). The black curves indicate the boundaries of the first significant time bins. (**B**) Same as (A) but for monkey C. (**C**) Topographic maps of maximum *F*-values within 300 ms post-stimulus for AC (left) and PFC (right) of monkey X. Channels with significant difference (< 350 ms) are circled in black and marked with ’o’. (**D**) Same as (C) but for monkey C.

**Supplementary Figure 6.** Topographic differential GC maps for all frequency bands at time window [0 100] ms. Channels with significant differences between responses to deviant and standard sounds are circled in black and marked with ‘o’ (one-tailed permutation test, *p* < 0.05, FDR corrected for channels).

**Supplementary Figure 7.** Averaged cross-period correlation matrices of differential GC maps between pairs in frequency ratios of 1.04, 1.06, and 1.08 for monkey. **C.** The color scale indicates Spearman’s ρ. Significant correlations for all pairs are indicated by dots for one monkey and by stars for both monkeys (Spearman’s rank correlation, *p* < 0.05, uncorrected).

**Supplementary Figure 8.** Information flow between AC and PFC in prediction under non-behavioral context. (**A**) Time-frequency maps of GC spectrum, aligned to the onset of the first standard sound and averaged across all channels for monkey X. The example GC was computed using trials with a standard sound number of 8. Traces on the right and left display the GC averaged across time from 1 to 3 s. (**B**) Topographic maps of GC correlation over time under 2-Hz and δ frequency bands for both monkeys. No channels with significant correlation were found.

**Supplementary Figure 9.** Averaged cross-period correlation matrices of differential GC maps between pairs in frequency ratios of 1.04, 1.06, and 1.08 under non-behavioral context. The color scale indicates Spearman’s ρ. Significant correlations for all pairs are indicated by dots for one monkey and by stars for both monkeys (Spearman’s rank correlation, *p* < 0.05, uncorrected).

## Acknowledgments

We are grateful to Xiaokai Kou for his help with the experiments. This work was supported by Brain Science and Brain-like Intelligence Technology–National Science and Technology Major Project (2022ZD0204600 and 2022ZD0204800) (to X.Y.); National Natural Science Foundation of China 32571216 and 32171044 (to X.Y.), and 32100827 (to Y.Z.); Project PID2023-148541OB-I00, funded by MICIU/AEI https://doi.org/10.13039/501100011033 and FEDER EU.; the Consejería de Educación, Junta de Castilla y León (SA218P23), and the strategic research programs of excellence from the Regional Government of Castile and León, co-funded by the ERDF Operational Programme (ref. CLU-2023-1-01), awarded to M.S.M.

