## Supplementary figures and images for "Behaviorally gated hierarchical predictive coding in macaque auditory–prefrontal circuits"

### Supplementary Figures 1-9

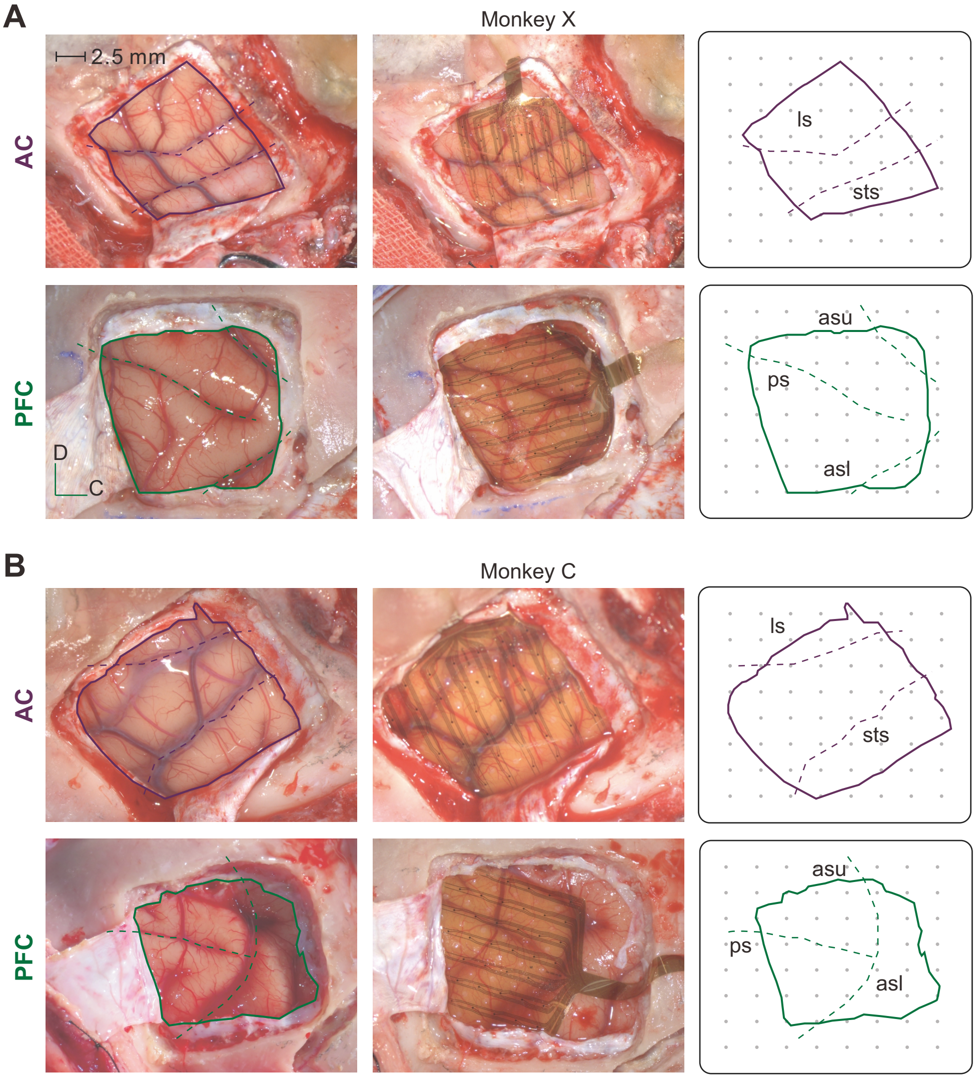

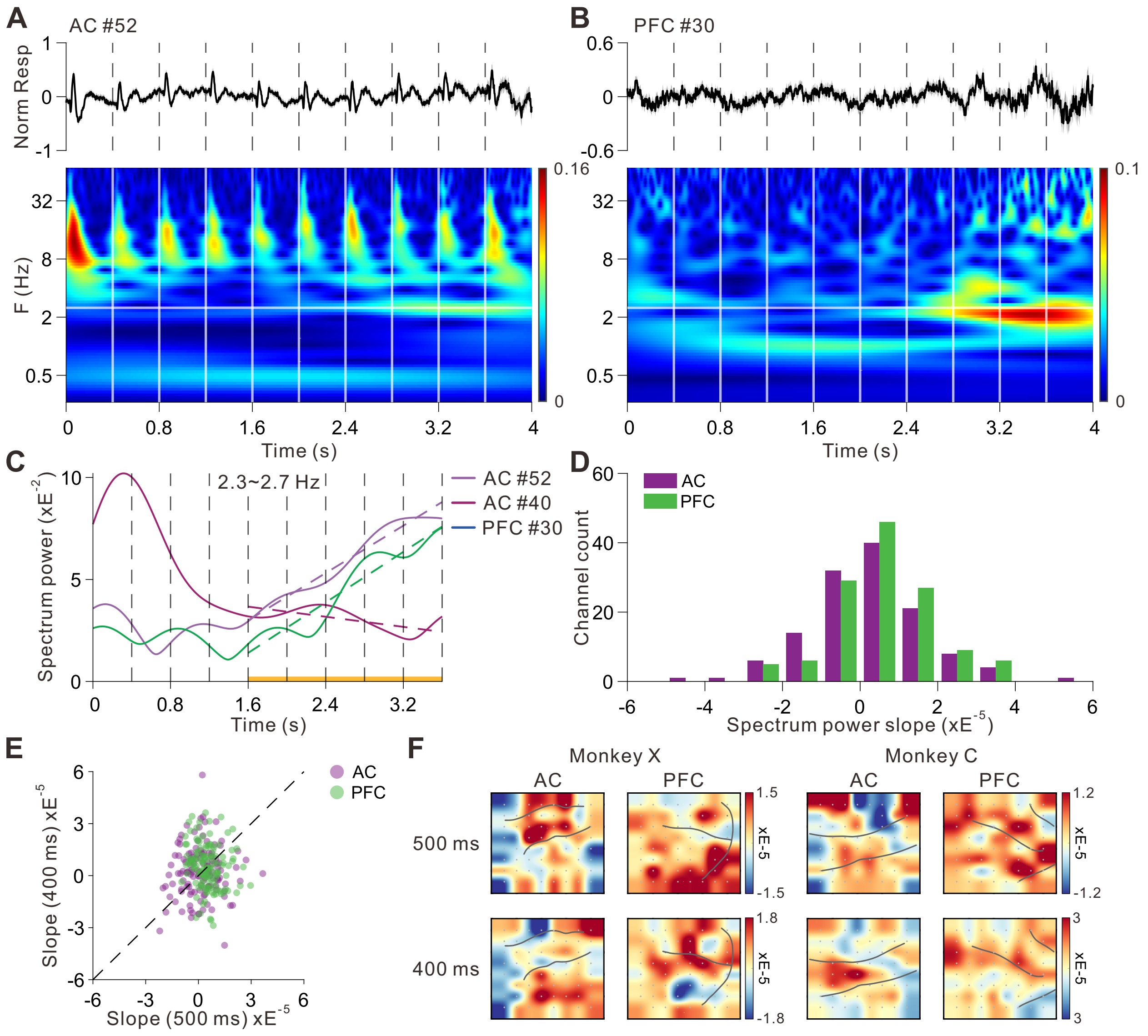

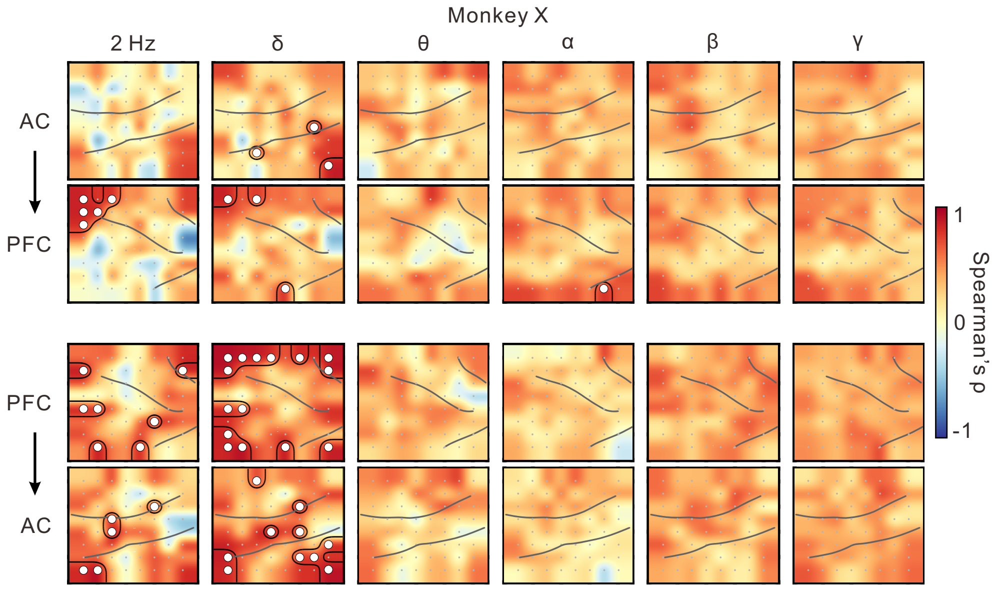

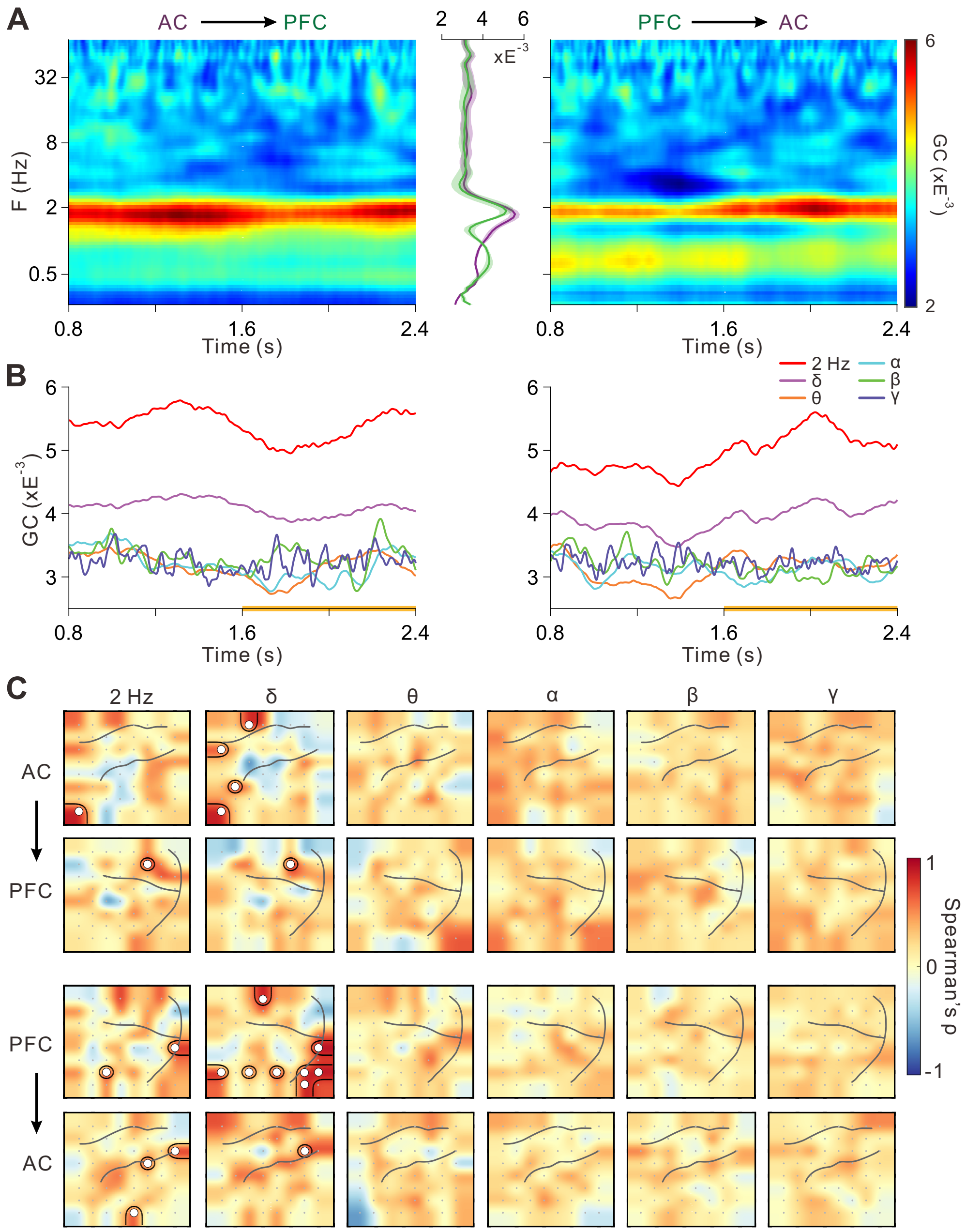

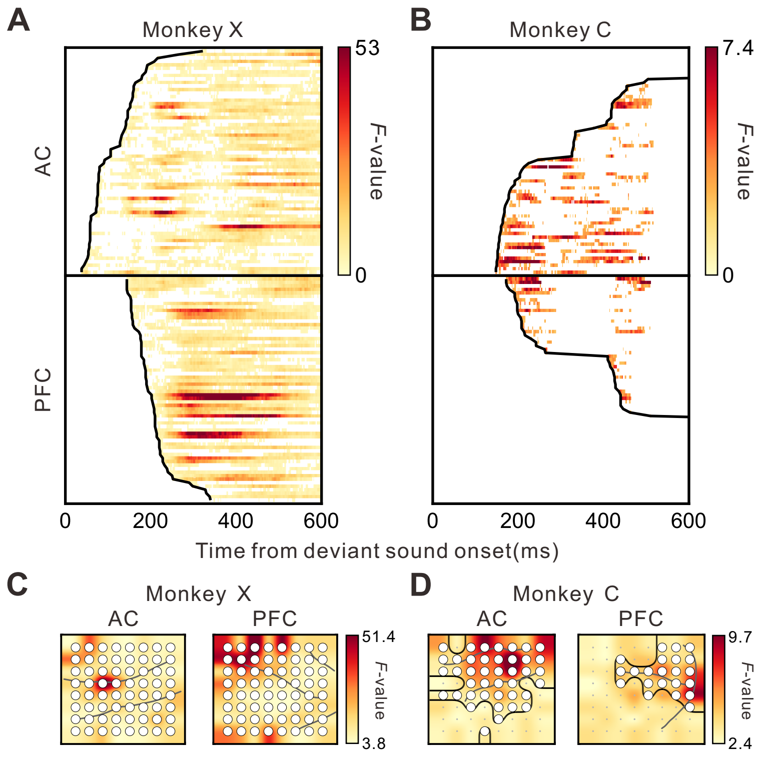

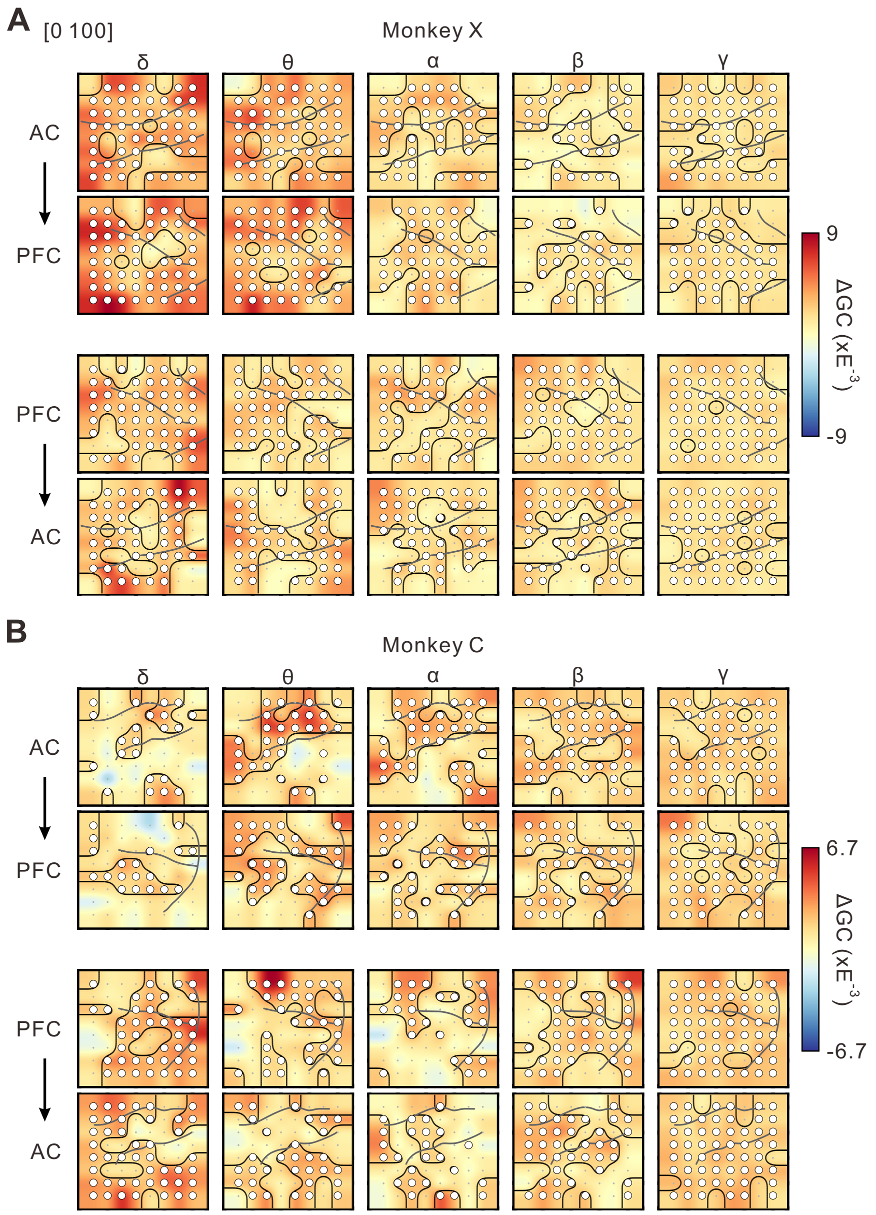

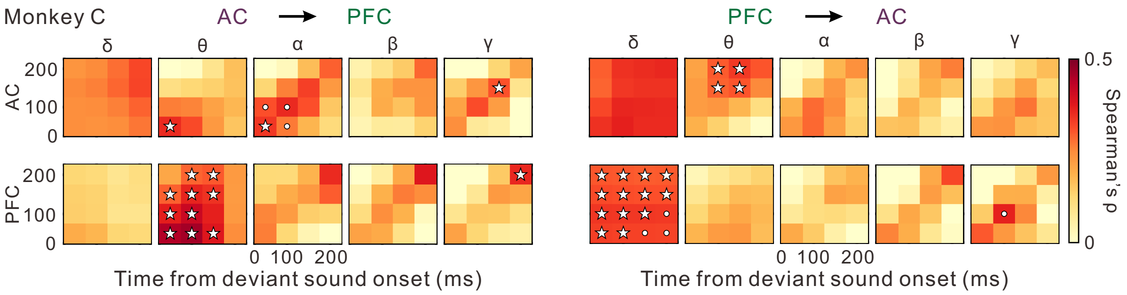

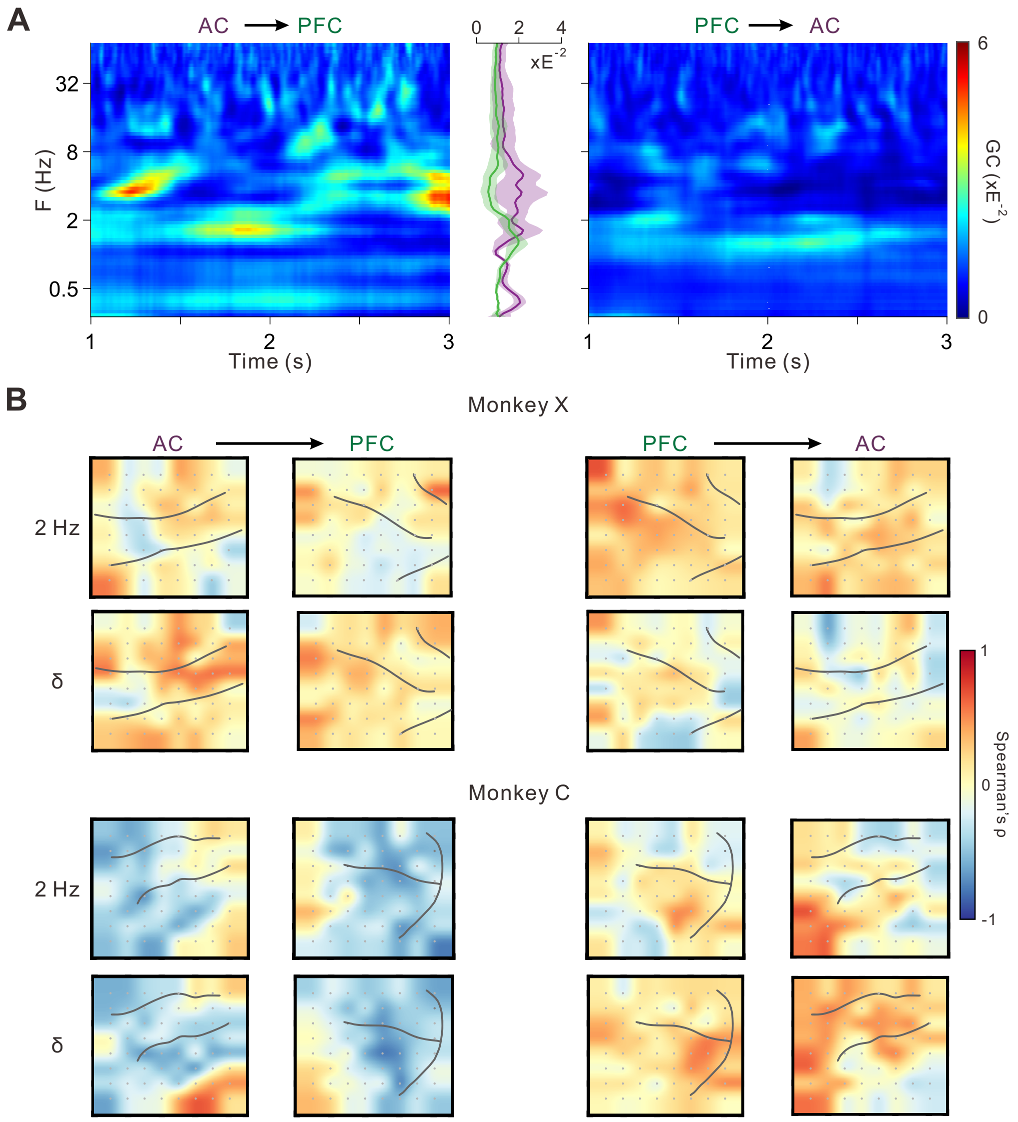

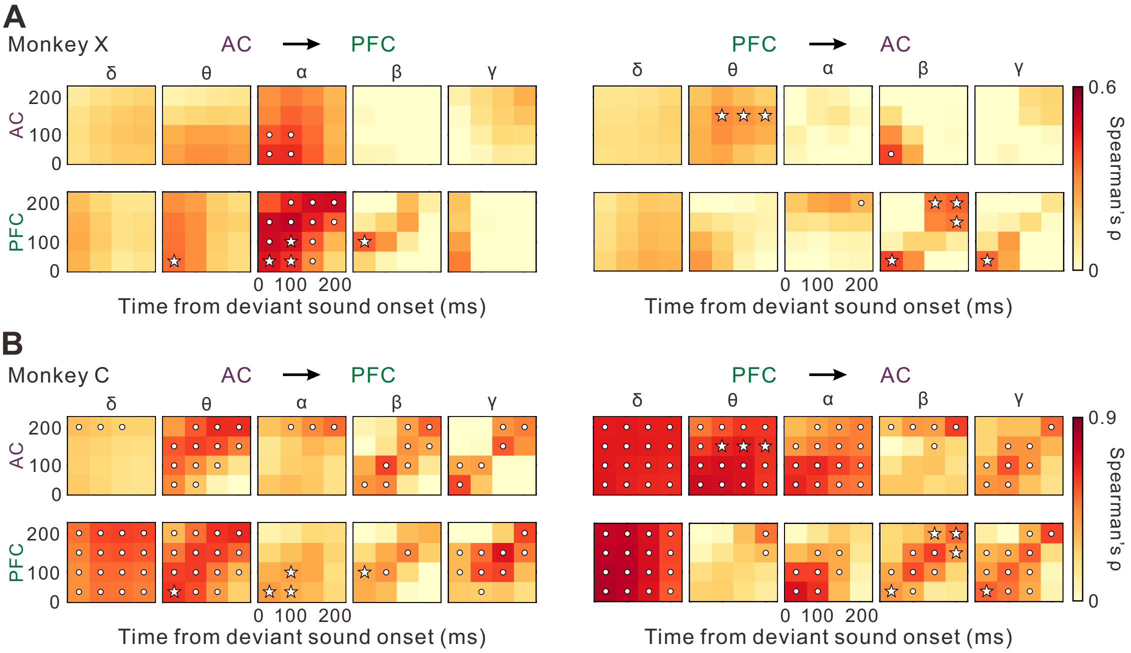
